# Cyclic gallium-68 labeled peptides for specific detection of human angiotensin-converting enzyme 2

**DOI:** 10.1101/2020.12.15.412809

**Authors:** Matthew F. L. Parker, Joseph Blecha, Oren Rosenberg, Michael Ohliger, Robert R. Flavell, David M. Wilson

**Author notes:** Correspondence and Reprint Request: David Wilson, M.D., Ph.D., Department of Radiology and Biomedical Imaging, University of California, San Francisco, 505 Parnassus Ave., San Francisco, CA 94143. First Author: Matthew F.L. Parker, Ph.D., Department of Radiology and Biomedical Imaging, University of California, San Francisco, 180 Berry St., San Francisco, CA 94107, Fax: (415).

## Abstract

In this study, we developed ACE2-specific, peptide-derived ^68^Ga-labeled radiotracers, motivated by the hypotheses that (1) ACE2 is an important determinant of SARS-CoV-2 susceptibility, and (2) that modulation of ACE2 in COVID-19 drives severe organ injury.

**Methods:** A series of NOTA-conjugated peptides derived from the known ACE2 inhibitor DX600 were synthesized, with variable linker identity. Since DX600 bears two cystine residues, both linear and cyclic peptides were studied. An ACE2 inhibition assay was used to identify lead compounds, which were labeled with ^68^Ga to generate peptide radiotracers ([^68^Ga]NOTA-PEP). The aminocaproate-derived radiotracer [^68^Ga]NOTA-PEP4 was subsequently studied in a humanized ACE2 (hACE2) transgenic model.

**Results:** Cyclic DX-600 derived peptides had markedly lower IC_50_’s than their linear counterparts. The three cyclic peptides with triglycine, aminocaproate, and polyethylene glycol linkers had calculated IC_50_’s similar to, or lower than the parent DX600 molecule. Peptides were readily labeled with ^68^Ga, and the biodistribution of [^68^Ga]NOTA-PEP4 was determined in a hACE2 transgenic murine cohort. Pharmacologic concentrations of co-administered NOTA-PEP (“blocking”) showed significant reduction of [^68^Ga]NOTA-PEP4 signals in the in the heart, liver, lungs, and small intestine. *Ex vivo* hACE2 activity in these organs was confirmed as a correlate to *in vivo* results.

**Conclusions:** NOTA-conjugated, cyclic peptides derived from the known ACE2 inhibitor DX600 retain their activity when N-conjugated for ^68^Ga chelation. *In vivo* studies in a transgenic hACE2 murine model using the lead tracer [^68^Ga]NOTA-PEP4 showed specific binding in the heart, liver, lungs and intestine - organs known to be affected in SARS-CoV-2 infection. These results suggest that [^68^Ga]NOTA-PEP4 could be used to detect organ-specific suppression of ACE2 in SARS-CoV-2 infected murine models and COVID-19 patients.

**TOC figure:** For Table of Contents use only

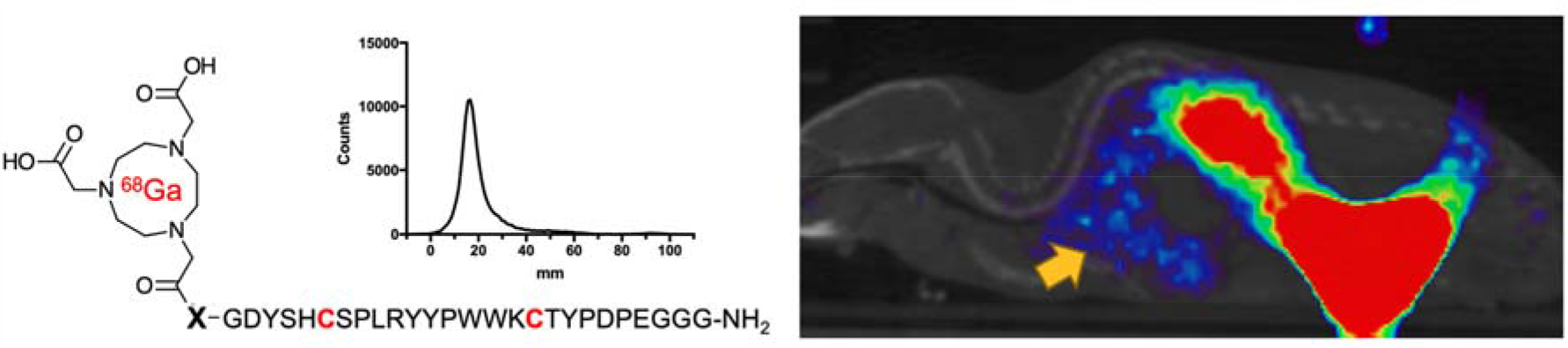

## Introduction

The novel coronavirus SARS-CoV-2 has had profound effects on global health, especially in the United States, the country with the largest number of confirmed COVID-19 cases, and associated deaths. Many of these patients progress to Acute Respiratory Distress Syndrome (ARDS), respiratory failure with widespread injury of the lungs. The underlying mechanisms include diffuse alveolar damage, surfactant dysfunction, and immune cell activation^1–3^. Of note, many pathologic conditions can cause this convergent picture, including both bacterial and viral infections. These causes of ARDS likely share dysfunction of the renin-angiotensin system, especially loss of angiotensin converting enzyme II (ACE2) function^4–8^. ACE2 is a transmembrane protein that functions as angiotensin receptor (ATR) chaperone. The roles of ACE2, ACE, and angiotensin II are highlighted in **Figure 1A** which describes dual functions of the renin-angiotensin system with opposing effects on cardiovascular biology^9^. In this pathway, ACE2 performs an important regulatory role, converting angiotensin II to angiotensin 1,7 which causes vasodilatation and has anti-inflammatory effect, unlike activation of ATR which will lead to vasoconstriction, higher blood pressure and inflammation (potentially ARDS)^10–13^.

**Figure 1.**
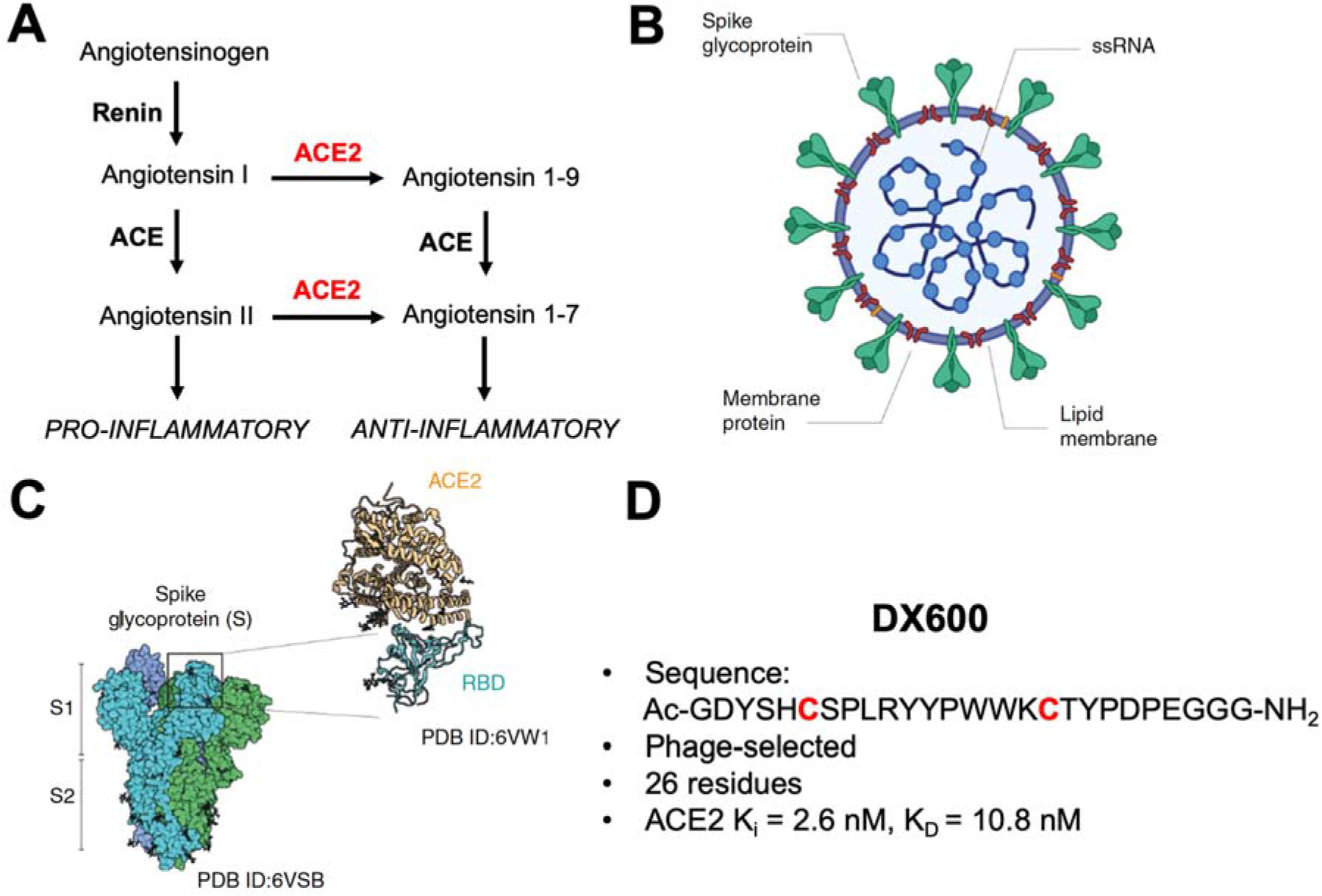
Role of ACE2 in hypertension and SARS-CoV-2 infection. (a) The renin-angiotensin system. The role of ACE2 highlighted on the right; ACE2 generally counters the “vasoconstrictive” pathway initiated by the formation of angiotensin II. (b) Simplified structure of the SARS-CoV-2 virus indicating the spike glycoprotein that interacts with ACE2 and other host proteins. (c) Structural (cryoEM, X-ray crystallography) and atomic force microscopy have elucidated the interaction between the spike protein S1 subunit and ACE2. Of note S1 binds to an ACE2 site remote to its active site, which is targeted by the inhibitory peptides described in this manuscript. Images adapted from Yang et al. 2020; source article is licensed under a Creative Commons Attribution 4.0 International License. (d) Characteristics of the 26-residue DX600 peptide. This peptide contains two cysteine residues, used for cyclization via disulfide bridge formation. DX600 was discovered via phage display and shown by Huang et al. to be a potent ACE2 inhibitor, a calculated K_D_ of 10.8 nM, and specificity for ACE2 versus angiotensin converting enzyme (ACE) and carboxypeptidase A (CPA).

Although several recent papers suggest that other mammalian transmembrane proteins (for example CD147 and CD26) allow SARS-C0V-2 to infect different cell types^14,15^, ACE2 is the main point of entry of the virus into host cells (**Figure 1B**). This process depends on this receptor as well as on its spike (S) protein, with cryo-EM and X-ray crystal structures of the complex recently described, as well as characterization of the complex via atomic force microscopy (**Figure 1C**)^16–18^. This protein has 2 subunits-S1 containing receptor binding domains (RBDs), and S2, which is responsible for membrane fusion. The RBDs can mimic the ACE2 interaction with ATR (hydrophobic and strong electrostatic interactions, including pi-pi, and cation-pi) and gain entry via strong non-covalent attachment to ACE2 in the ATR binding site^19^. Three recent cryo-EM studies demonstrated that SARS-CoV-2 spike protein directly binds to ACE2 and that the SARS-CoV-2 spike protein likely recognizes human ACE2 with even higher binding affinity than spike from SARS-CoV^20–22^. This binding was suggested to alter virus configuration and expose a cleavage site on S2, resulting in host protease cleavage (mainly by transmembrane protease/serine subfamily member 2 - TMPRSS2), allowing the virus to enter the cell^23^. This mechanism was recently supported by a cryo-EM post fusion analysis that showed structural and conformational rearrangements of the S-protein compared to its pre-fusion structure^24^.

To investigate SARS-CoV-2 susceptibility, and organ-specific suppression of ACE2 in COVID-19, new ACE2-specific imaging methods would be profoundly helpful. However, ACE2-specific small molecules and peptides developed following the original 2003 SARS-CoV outbreak (this virus also depends on ACE2 for viral entry) offer clues as to how active-site targeted, high affinity ligands might be developed. A large number of ACE2-specific ligands have been reported, generally characterized by their ACE2 IC_50_’s, and including the peptide DX600 discovered via phage display^25–30^. The DX600 sequence is shown in **Figure 1D**. In this manuscript, we report development of ACE2-specific PET radiotracers ([^68^Ga]NOTA-PEP) derived from this sequence. We anticipate that ACE2-specific PET could help evaluate which systems are most targeted by SARS-CoV infection, the timing of disease, and how ACE2 modulation correlates with ARDS susceptibility and other organ injury. Recent work has highlighted the role of ACE2 in a large number of organs beyond the lungs, including the heart, kidneys, and gastrointestinal system^31–36^. We therefore believe that the information gleaned from [^68^Ga]NOTA-PEP4 or some other *in vivo* ACE2 sensor will potentially be helpful in COVID-19 treatment, either via exogenous ACE2^4,37^ or some other therapy.

## Materials and Methods

### Peptides

The DX600-derived peptides studied were obtained from AnaSpec (Fremont, CA) as a custom synthesis, fully characterized by HPLC and mass spectrometry. These peptides were radiolabeled without additional modification.

### ACE2 inhibition assay

Six DX600-derived peptides, named NOTA-PEP1-6, (cyclic versus non-cyclic, with triglycine, aminocaproate, and polyethylene glycol linkers) were studied using a commercially available ACE2 inhibition assay according to the manufacturer’s instructions (SensoLyte® 390 ACE2 Activity Assay Kit *Fluorimetric*, AS-72086, AnaSpec, Fremont CA). Each peptide inhibitor was first tested at 4 concentrations. Initial velocities were determined relative to the inhibitor free reaction. Subsequently, IC_50_ values were derived from nonlinear fits of saturation curves of a 6-point dilution series of peptide inhibitors.

### [^68^Ga] peptide synthesis

Full descriptions of radiochemical syntheses, as well as the analytical techniques used, are provided in the Supplemental Information. Unless otherwise noted, all reagents were obtained commercially and used without further purification. [^68^Ga]gallium chloride was generated in the UCSF radiopharmaceutical facility by elution from an ITG germanium-gallium generator. To generator eluted [^68^Ga]Cl_3_ in a 4mL dilute HCl solution was added the indicated NOTA-PEP precursor (80µg) in pH 5 sodium acetate buffer solution (160µL). The mixture was heated for 15 mins at 90°C. The reaction was monitored by TLC performed on cellulose filter paper developed in PBS. Free gallium migrates to the solvent front (∼90 mm) and bound gallium remains at the origin (∼20 mm).

### μPET/CT imaging

All animal procedures were approved by the UCSF Institutional Animal Care and Use Committee, and all studies were performed in accordance with UCSF guidelines regarding animal housing, pain management, and euthanasia. Humanized ACE2 recombinant mice (B6.Cg-Tg(K18-ACE2)2Prlmn/J, 034860, Jackson Laboratory) were obtained from Jackson Labs, aged 6-10 weeks^38–40^. *Single time-point imaging:* A tail vein catheter was placed in mice under isoflurane anesthesia. Approximately 350 μCi of [^68^Ga]NOTA-PEP4 were injected via the tail vein catheter. The animals were placed on a heating pad to minimize shivering. Mice were allowed to recover, micturate, and at 75 minutes post-injection, placed back under isoflurane anesthesia. At 90 mins post-injection, the animals were transferred to a Siemens Inveon micro PET-CT system (Siemens, Erlangen, Germany), and imaged using a single static 25 min PET acquisition followed by a 10 min micro-CT scan for attenuation correction and anatomical co-registration. No adverse events were observed during or after injection of any compound. Anesthesia was maintained during imaging using isofluorane. *Inhibition (“blocking”) studies:* The protocol was identical to that above but cold NOTA-derived inhibitory cyclic peptide (NOTA-PEP4) (10mg/mL concentration) was co-administered with [^68^Ga]NOTA-PEP4 in buffered saline. *Dynamic imaging:* The protocol was similar to above except tail-vein administration of 350 μCi of [^68^Ga]NOTA-PEP4 was performed simultaneously on a cohort of 4 animals in bed positioning for PET imaging. PET imaging data was collected beginning at the moment of injection for 90 minutes followed by 10-minute CT.

### *Ex vivo* analyses of mice

Upon completion of imaging, mice were sacrificed and biodistribution analysis performed. Gamma counting of harvested tissues was performed using a Hidex Automatic Gamma Counter (Turku, Finland). Organs were also harvested from a separate cohort of mice for an ACE2 activity assay. The tissues were homogenized and aliquots were used for protein concentration using a standard Bradford assay. Additional tissue aliquots were used as the source of ACE2 in a commercially available ACE2 assay (AnaSpec, Fremont, CA). The initial velocities were normalized relative to muscle tissue. Relative activities are reported as the relative initial velocity/g of protein.

### Data analysis and statistical considerations

For syntheses, radiochemical yields incorporate decay-correction for ^68^Ga (t_1/2_=68 min). All *in vivo* PET data were viewed using open source Amide software (amide.sourceforge.net). Reported static (single time-point data) reflects gamma counting of harvested tissues. For dynamic data, quantification of uptake was performed by drawing spherical regions of interest (5-8 mm^3^) over indicated organs on the CT portion of the exam, and expressed as percent injected dose per gram. All statistical analysis was performed using Microsoft Excel. Data were analyzed using an unpaired two-tailed Student’s t-test. All graphs are depicted with error bars corresponding to the standard error of the mean.

## Results

### NOTA-conjugated, cyclic peptides targeting the ACE2 active site retain their potency relative to the DX600 parent compound

Based on our hypothesis that potent peptide-derived ACE2 inhibitors, modified with linkers/chelating groups will retain their activity and specificity, several NOTA-modified peptide-derived ACE2 inhibitors derived from the DX600 sequence^29^ (K_i_ = 2.8 nM, K_d =_ 10.8 nM) were synthesized and screened for ACE2 inhibition. These were synthesized via Fmoc-protected linkers and N-capping NOTA reagents (**Figure 2A**,**B**). The general structure pursued was a NOTA-linker-peptide with three different linkers used, conferring varying degrees of hydrophobicity and hydrogen bonding: triglycine, PEG, or caproic acid. These were synthesized using standard Fmoc solid-phase synthesis^41^ (AnaSpec, Fremont CA) with purity and identity confirmed by HPLC and mass spectrometry. Because DX600 contains two cysteine residues, a cyclized set of peptides were also synthesized via disulfide bridge formation^42^. When these compounds were compared to the parent DX600 peptide in a commercially available fluorometric ACE2 inhibition assay (AnaSpec), all three cyclic peptides (NOTA-PEP2, NOTA-PEP4, NOTA-PEP6) showed ACE2 inhibition nearly identical to DX600 (**Figure 2C**,**D**,**E**). In other words, the N-terminal modification caused no loss of inhibitory activity when compared to the parent peptide, and in fact the cyclic peptide NOTA-PEP4 was a slightly better ACE2 inhibitor than DX600. In contrast, the linear derivatives showed much lower activity, which may result from a solution confirmation for which the NOTA interferes with ACE2 active site binding. To further evaluate this loss of potency, we studied ACE2 inhibition using a cyclic NOTA-PEP with and without addition of the reducing agent tris(2-carboxyethyl)phosphine (TCEP) which is expected to reduce the disulfide bridge in the cyclic peptide (producing the linear NOTA-PEP5) (**Figure 2F. Supp. Fig. 1**). As anticipated, addition of TCEP markedly increased the observed ACE2 IC_50_.

**Figure 2.**
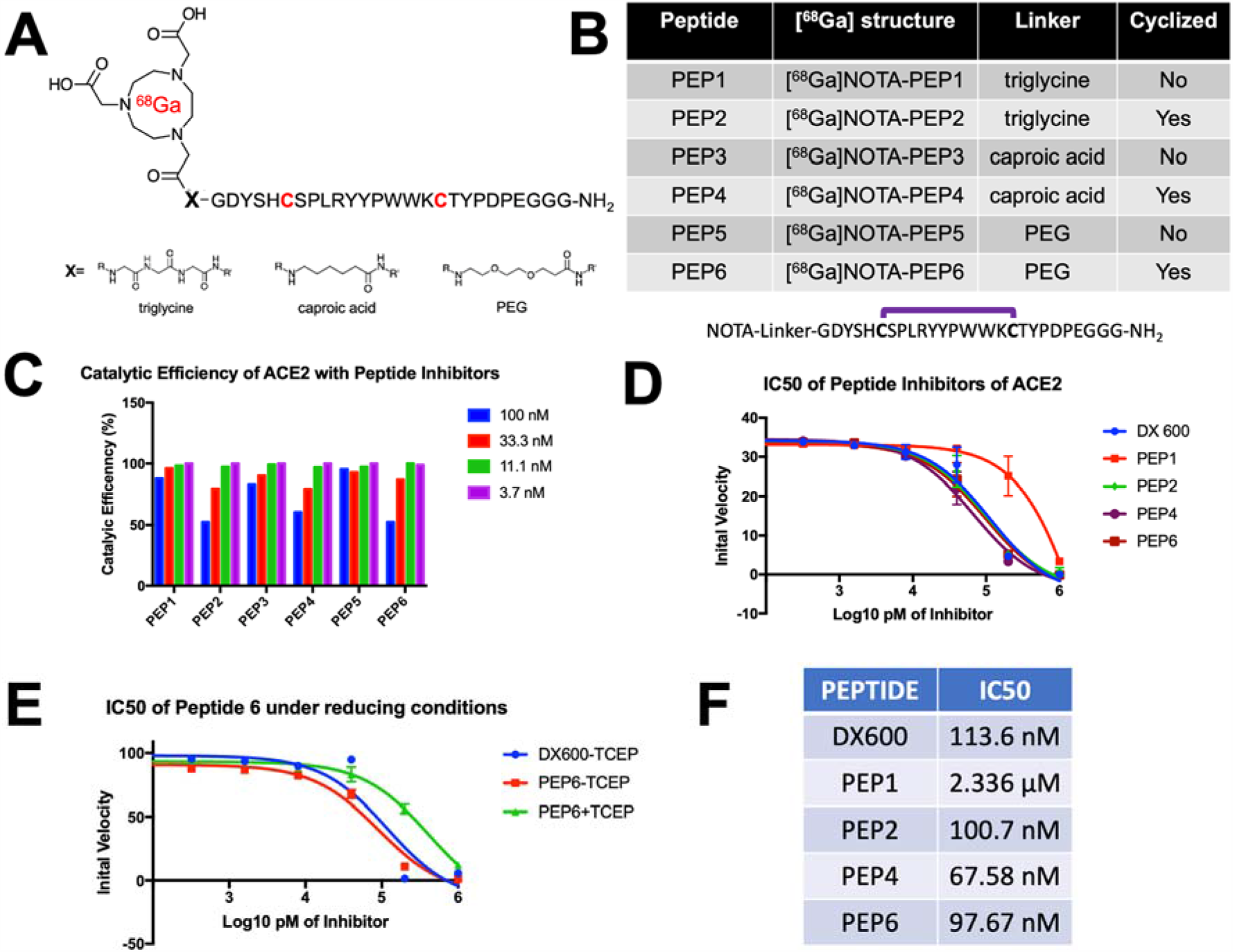
Discovery of DX600-derived, NOTA-conjugated cyclic peptide inhibitors of ACE2 from a small library. (a) General [^68^Ga] peptide structure pursued. Peptides studied had an N-terminal NOTA chelating group, triglycine/caproic acid/PEG linkers with varying degrees of hydrophobicity and hydrogen-bonding, and +/- cyclization via the cysteine residues highlighted in red. (b) Identity of 6 NOTA-conjugated peptides studied. (c) ACE2 catalytic efficiency in the presence of DX600-derived inhibitors. Cyclic peptides were markedly more potent than their linear counterparts. (d) Greater potency of the cyclic peptides are highlighted by the initial ACE2 velocities seen with increasing inhibitor concentrations. All cyclic peptides (NOTA-PEP2, NOTA-PEP4, NOTA-PEP6) had similar profiles to the parent peptide DX600, in contrast to linear peptide NOTA-PEP1. (e) The effects of cyclization were highlighted in a separate ACE2 assay using TCEP to reduce the disulfide bridges in NOTA-PEP6. (f) ACE2 IC_50_’s calculated from these data. Of note these IC_50_’s are significantly higher than the K_i_’s reported by Huang et al. for DX600, likely reflecting differences in the assays used. However, most importantly NOTA-conjugated cyclic derivatives had no loss of potency relative to the DX600 parent.

### Efficient radiosyntheses of [^68^Ga]NOTA-PEP peptides

Promising ACE2 inhibition results for NOTA-conjugated cyclic peptides were followed with radiolabeling of peptides with ^68^Ga ^43^ (**Supp. Fig. 2**). Crude radiochemical purities of the desired [^68^Ga]-peptide chelate were > 90% in all cases. The majority of synthetic efforts focused on optimizing the radiosynthesis of the lead inhibitor [^68^Ga]NOTA-PEP4. [^68^Ga]NOTA-PEP4 was synthesized in 30mins from generator eluted [^68^Ga]Cl_3_ in a 4mL dilute HCl solution. The precursor (80µg) was added as a pH 5 acetate buffer solution (160µL) and heated for 15 mins at 90°C. Radio TLC (**Figure 3A**) was performed on cellulose filter paper developed in PBS. The final radiochemical purity of [^68^Ga]NOTA-PEP4 was > 95% in all cases with a decay-adjusted radiochemical yield of 63.2 ± 6.4% (N = 8).

**Figure 3.**
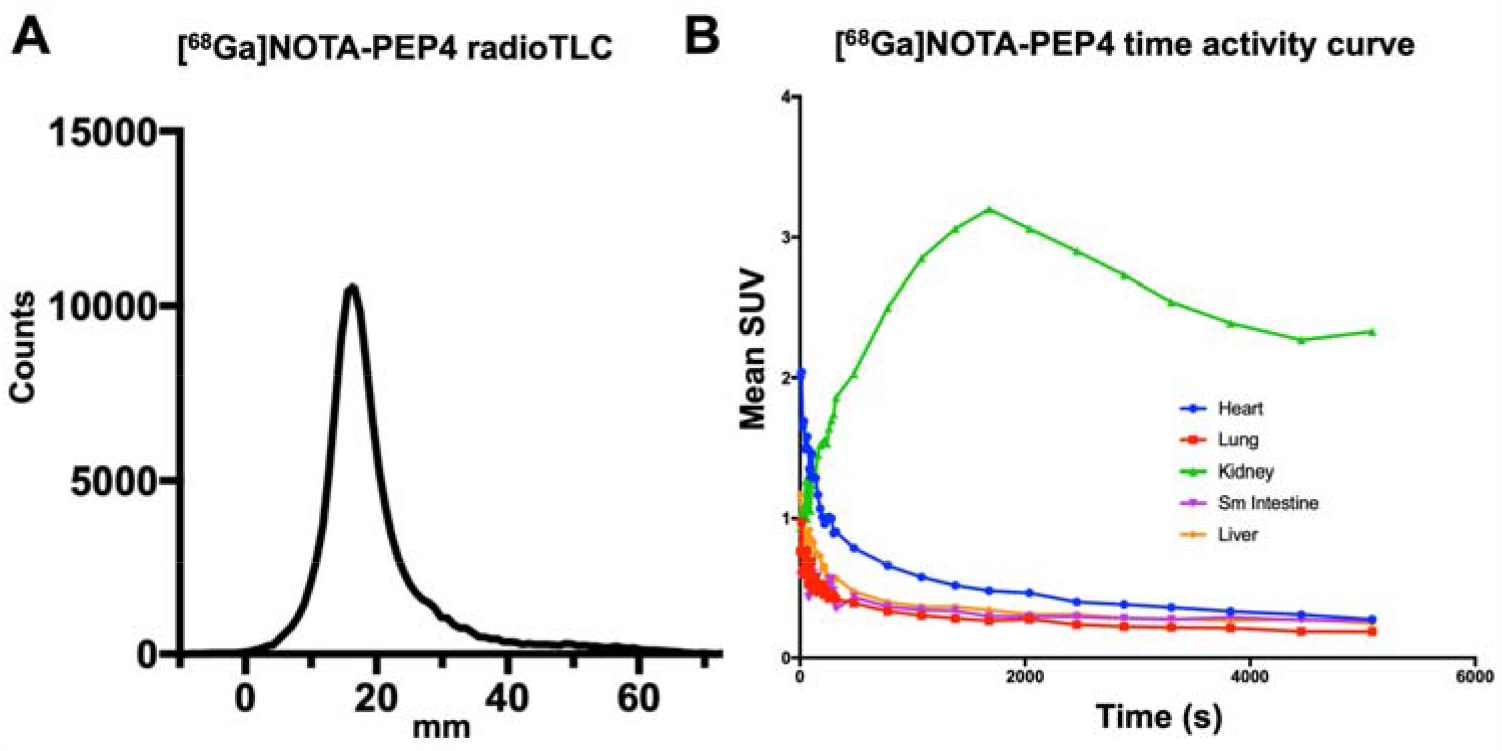
Radiosynthesis and *in vivo* dynamic characterization of [^68^Ga]NOTA-PEP4. Based on IC_50_ data, NOTA-PEP4 was chosen for subsequent radiolabeling with ^68^Ga. (a) Our optimized radiosynthesis yielded the desired [^68^Ga]NOTA-PEP4 in greater than 95% radiochemical purity. (b) Dynamic μPET-CT in hACE2 transgenic mice was used to generate an organ-specific time-activity curve, identifying later time points as generating stable [^68^Ga] signals.

### [^68^Ga]NOTA-PEP4 signals in the lungs, heart, small intestine and liver of hACE2 transgenic mice are attenuated with co-administration of inhibitory cyclic peptide

Having developed a radiosynthesis of [^68^Ga]NOTA-PEP4, we sought to further validate the tracer in a transgenic, humanized ACE2 (hACE2) murine model. The *K18-hACE2* transgenic mice express human ACE2 under the control of the human keratin 18 promoter, which directs expression to epithelia, including airway epithelial cells where infections typically begin^38^. Preliminary studies available from the Jackson Laboratory website, and recently published studies^44^ have shown that *K18-hACE2* transgenic mice develop dose-dependent disease phenotypes when infected intranasally with SARS-CoV-2 with high doses resulting in ARDS/death analogous to that observed in some COVID-19 patients. Male Tg(K18-ACE2)2Prlmn/J hemizygous mice (N = 4, Jackson Lab) were initially injected with 13.0 MBq (350 µCi) of [^68^Ga]NOTA-PEP4 and dynamic imaging was performed to identify optimum single time-point imaging. Region of interest (ROI) analysis of dynamic data was focused on organs known to be affected in SARS-CoV-2 (**Figure 3B, Supp. Fig. 3**). ROI analysis of the images demonstrated prompt clearance from the blood pool with accumulation in the kidneys, as expected for a small peptide tracer.

Next, we performed an imaging and biodistribution study, to show that [^68^Ga]NOTA-PEP4 demonstrates specific uptake in tissues with increased expression of ACE2. In order to demonstrate specificity of uptake, blocking with excess cyclic NOTA-PEP inhibitory peptide was employed. With blocking, significant reductions in cyclic [^68^Ga]NOTA-PEP4 were seen in the heart (p = 0.0203), lung (p <0.0001), liver (p <0.0001) and small intestine (p = 0.0002). ACE2 activity in these organs was subsequently confirmed via harvested organs in a separate hACE2 cohort (N = 3, **Supp. Fig. 4**). Taken together, these data demonstrate that [^68^Ga]NOTA-PEP4 can specifically bind to tissues with high ACE2 expression.

**Figure 4.**
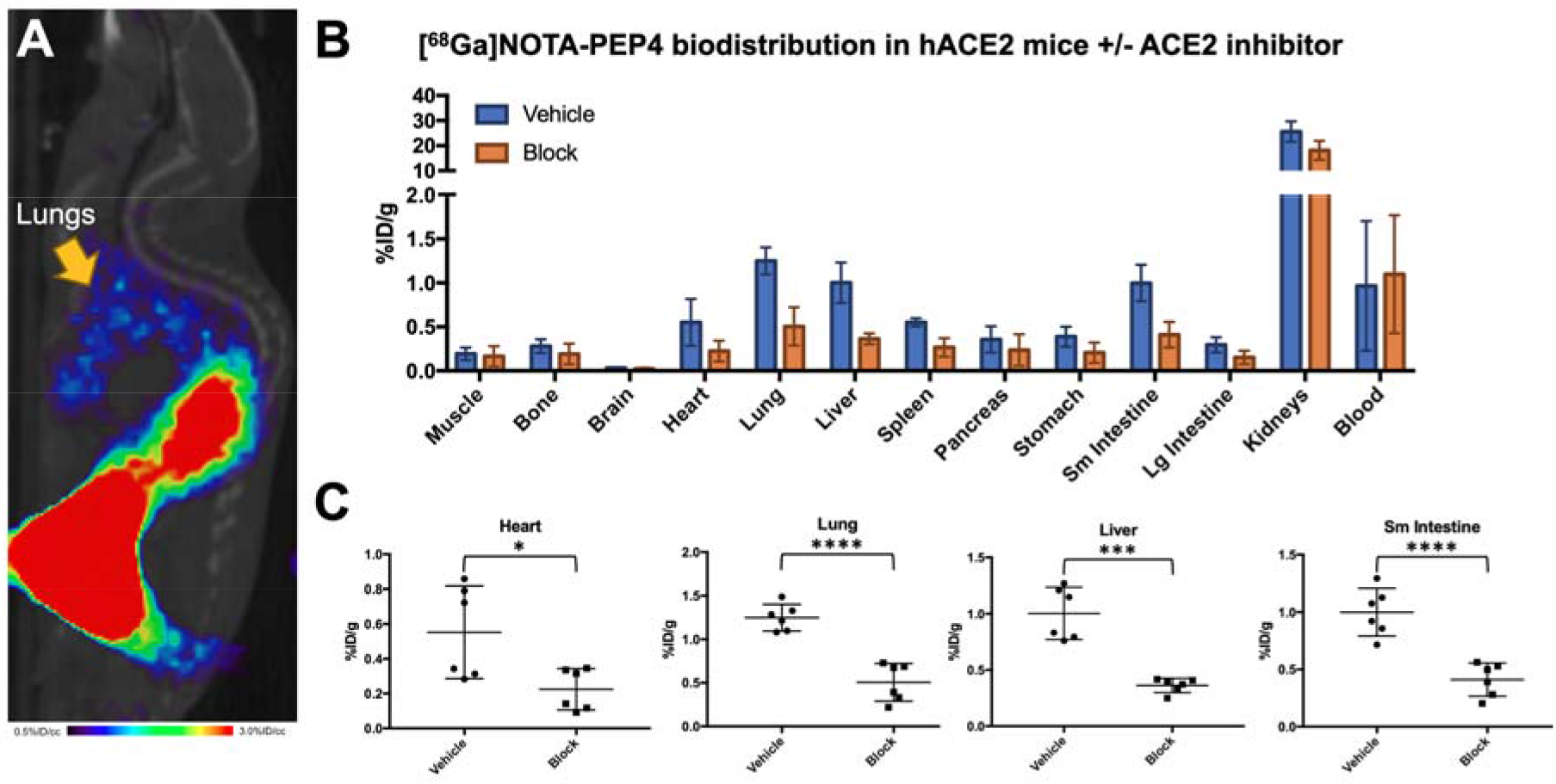
*In vivo* biodistribution studies of [^68^Ga]NOTA-PEP4 in hACE2 transgenic mice, demonstrating modulation of signals with a pharmacologic dose of ACE2 inhibitor. (a) μPET-CT image from a static acquisition highlighting the signal corresponding to the lungs, which are of exceptional interest in SARS-CoV-2 infection. (b) Biodistribution studies with and without the presence of an ACE2 inhibitor. The highest signals were observed in the kidneys, but the observed %ID/g was not significantly lower in the presence of ACE2 inhibitor. Therefore renal signals are felt to represent the primary route of excretion. (c) Significant blocking of [^68^Ga]NOTA-PEP4 was seen in the heart, lungs, liver, and small intestine, organs implicated in COVID-19.

## Discussion

The novel coronavirus disease (COVID-19) has spread rapidly throughout the world with the highest number of confirmed cases and deaths in the United States. Both biochemical studies and published cryo-EM structures have shown that the spike protein (S-protein) of severe acute respiratory syndrome coronavirus 2 (SARS-CoV-2) predominantly uses human angiotensin-converting enzyme 2 (ACE2) for viral entry, resulting in suppression of this enzyme as seen in SARS-CoV^45,46^. Additional recent publications have highlighted the possibility that the lower ACE2 activity seen with SARS-CoV-2 infection may be responsible for the physiologic effects incurred, analogous to what was seen with the original SARS-CoV^46^. These observations support recombinant ACE2-derived therapies as a way to treat COVID-19, via two mechanisms: (1) by replenishing “protective” ACE2 function and (2) by serving as a “decoy” receptor for the virus. These therapeutic effects, the differential susceptibility of individuals (based on age, co-morbidities) to COVID-19, and the organ-specific effects of SARS-CoV-2 are all potentially addressed by an ACE2-specific imaging method. We therefore sought a PET tracer derived from known inhibitor structures, via modification of the known ACE2 inhibitory peptide DX600 with ^68^Ga.

Inhibitor-derived structures modified for PET do not necessarily recapitulate the potency of their parent compounds, so our first efforts focused on the “cold” NOTA-conjugated DX600 derived peptides, derived from triglycine, caproic acid, and PEG linkers. Gratifyingly, the DX600-derived cyclic peptides studied all showed ACE2 activity similar to the parent peptide. In contrast, the linear versions were relatively inactive, which may reflect conformational effects. Of note, the calculated IC_50_ of DX600 (standard included in AnaSpec assay kit) was > 1 order of magnitude higher than the K_i_reported by Huang et al.^29^, likely reflecting numerous experimental differences (enzyme concentration and activity, etc.). We therefore considered the IC_50_ of the NOTA-derived peptides relative to that of DX600 to be the most important determinant of successful PET probe development. Indeed, our lead cyclic peptide NOTA-PEP4 had an IC_50_ lower than that of the DX600 parent, motivating the radiolabeling of NOTA-PEP4 for subsequent imaging studies.

A high-yield and efficient synthesis of [^68^Ga]NOTA-PEP4 was developed with the tracer applied to a hACE2 transgenic model. Our studies co-injecting a pharmacologic concentration of NOTA-PEP inhibitor with [^68^Ga]NOTA-PEP4 showed significant attenuation of PET signals in the lungs, liver, heart, and small intestine-suggesting that these signals were related to ACE2 expression. Consistent with this observation, *ex vivo* tissue-specific ACE2 activity was observed in these organs, which are affected in COVID-19^47,48^. Modulation of [^68^Ga]NOTA-PEP4 using an ACE2 inhibitor also suggests that changes in ACE2 expression can be detected non-invasively. Additionally, *ex vivo* tissue analysis showed metabolically activity ACE2 expression in the kidneys despite the absence of strong “blocking.” The tissue accumulation of [^68^Ga]NOTA-PEP4 in the kidneys suggests a dominant renal excretion pathway, complicating our ability to detect hACE2 in this tissue^49^. In other words, high background signal due to the normal excretion pathway of [^68^Ga]NOTA-PEP4 may represent a limitation of this method to detect ACE2 activity in the kidney. In the future, hACE2 expression-specific [^68^Ga]NOTA-PEP4 signals versus background excretion needs to be further clarified, perhaps using koACE2 animals^50^ in addition to the inhibitory studies described in this manuscript.

The *in vivo* studies performed also reflect a limitation of most academic centers in the United States; specifically, few facilities have a biosafety level 3 (BSL-3) compatible μPET-CT imaging system. Future molecular imaging of live SARS-CoV-2 (a BSL-3 organism) and its host effects will therefore require collaborative work with those few centers able to accommodate these studies^51^. Given the history of ACE2 with respect to SARS-CoV (the 2003 SARS coronavirus) and ARDS, we expect that new ACE2-specific PET tools will be relevant beyond the current pandemic. We are partially motivated by data indicating that zoonotic infections especially coronavirus-related are on the rise^52^. The incidence of emerging and re-emerging zoonotic disease is increasing in many parts of the world, with animal viruses able to cross species barriers to infect humans; it appears likely that ACE2 will be relevant in future pandemics. Better understanding ACE2 suppression, and differential susceptibility to SARS-COV-2 will help us better treat COVID-19 and other diseases for which ACE2 plays a critical role.

## Conclusions

Our study shows that the ACE2 active site-targeted inhibitor DX600 can be modified for PET via NOTA/linker modification, without loss of activity for cyclized peptides. All peptides studied are readily radiolabeled with ^68^Ga. In a humanized ACE2 transgenic murine model, the lead radiotracer [^68^Ga]NOTA-PEP4 shows dominant excretion from the kidneys, with attenuated uptake in the lungs, liver, heart, and small intestine when an ACE2 inhibitor is co-administered. These results suggest that modulation of ACE2, as occurring in SARS-CoV-2 infection, can be detected using [^68^Ga]NOTA-PEP4 or related approaches. Future studies include application of [^68^Ga]NOTA-PEP4 to SARS-CoV-2 infected hACE2 mice.

## Author contributions

DMW proposed and supervised the overall project. MP, JB, RF performed or supported the radiochemistry. MP performed μPET-CT imaging studies and subsequent data analysis. MP performed *ex vivo* analysis. DMW, MP, JB, RF, OR, MO wrote and edited the paper.

### Corresponding Author

*

### Notes

The authors declare no competing financial or other interests.

## Acknowledgements

Grant sponsors NIH R01 EB024014; R01 EB025985; UCSF Resource Allocation Program. The authors would also like to thank Prof. Sanjay Jain and Alvaro Ordonez (Johns Hopkins University) for helpful conversations.

## Supplementary information

Please see the supplementary information for detailed information regarding synthesis, and several imaging studies not reported in the main text.

