## Supplementary material for "Cyclic gallium-68 labeled peptides for specific detection of human angiotensin-converting enzyme 2": Supp

San Francisco, CA 94158, USA

[2]Department of Medicine

University of California, San Francisco

San Francisco, CA 94158, USA

[3]Department of Radiology

Zuckerberg San Francisco General Hospital

San Francisco CA 94110, USA

---

**D. AnaSpec peptide QC Data**……………………………………………………………………………..4

#### A. Determination of ACE2 $IC_{50}$'s:

| PEPTIDE | IC50 |
|---|---|
| DX600 | 118.2 nM |
| 6-TCEP | 90.93 nM |
| 6+TCEP | 412.5 nM |

**Supp. Fig. 1:** ACE2 $IC_{50}$'s for NOTA-ACE2pep6 when TCEP is used for disulfide bridge reduction (conversion of cyclic to linear peptide). An approximately 5-fold higher calculated $IC_{50}$ was observed in the presence of TCEP.

#### B. Radiochemistry:

80 µg of NOTA-peptide (2 µg/µL concentration) was diluted into 160 µL of sodium acetate buffer (pH = 5.5) and added to $^{68}GaCl3$ solution (370-740 MBq in 4 mL 0.05 M HCl; eluted from a generator). The mixture was tested for pH and additional sodium acetate buffer (pH = 5.5) was added to modulate the pH to between 3.5-4. The mixture was heated to 90°C for 10 minutes, diluted with 50 mM ammonium acetate solution and loaded onto a preconditioned C18 Sep-Pak. The cartridge was washed with 50 mM ammonium acetate solution and the labelled peptide was eluted from the cartridge in 200 µL fractions with 70:30 ethanol/50 mM ammonium acetate solution. The most concentrated fraction was diluted 10-fold with 0.9% sodium chloride solution and passed through a sterile filter for use. [$^{68}$Ga]-NOTA-ACE2pep4 was obtained 63% yield (decay corrected, n = 8) in greater than 95% purity (n = 8).

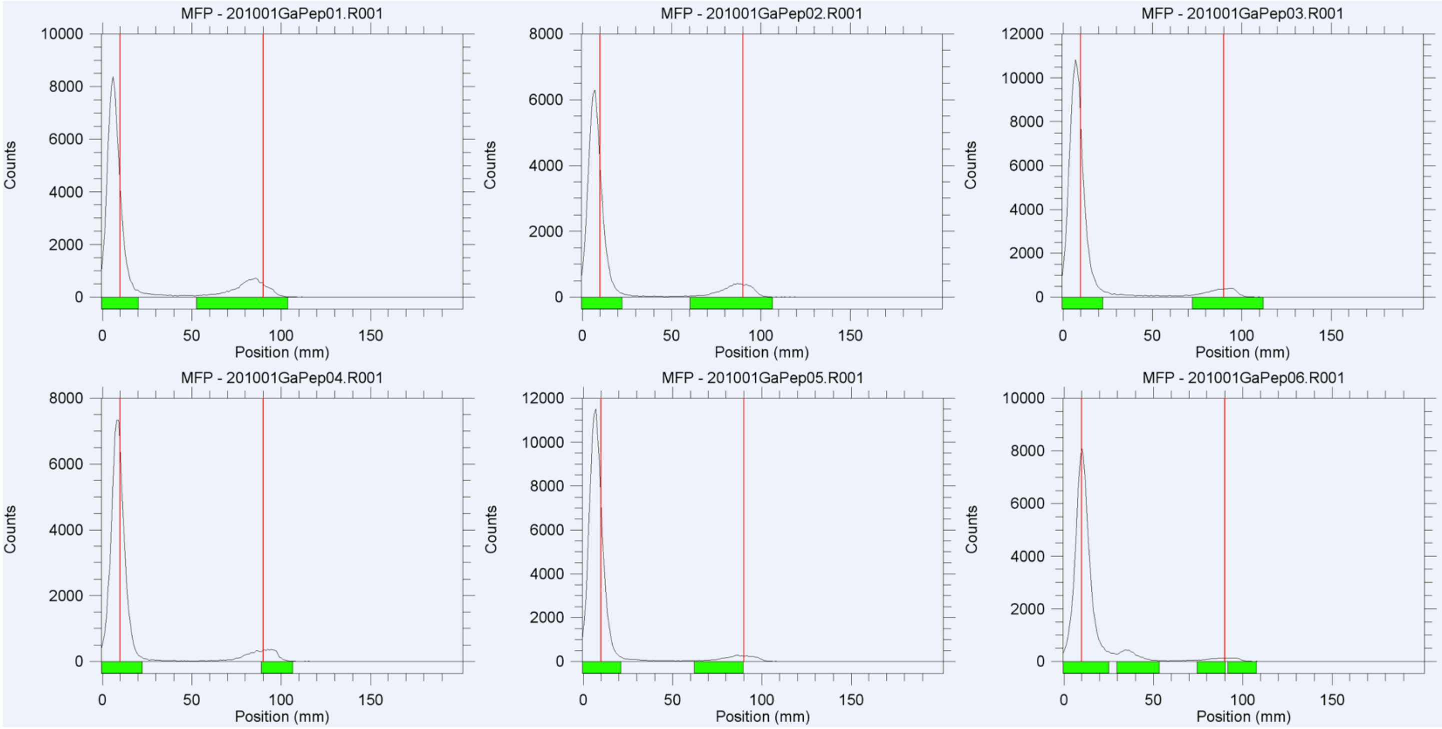

**Supp. Fig. 2:** Radiosyntheses of all six [$^{68}$Ga]-NOTA-ACE2pep peptides (crude RadioTLC). The chelated [$^{68}$Ga]-NOTA-ACE2pep was > 90% of radioactive signals observed in the crude product, in all cases.

#### C. Supplemental Figures:

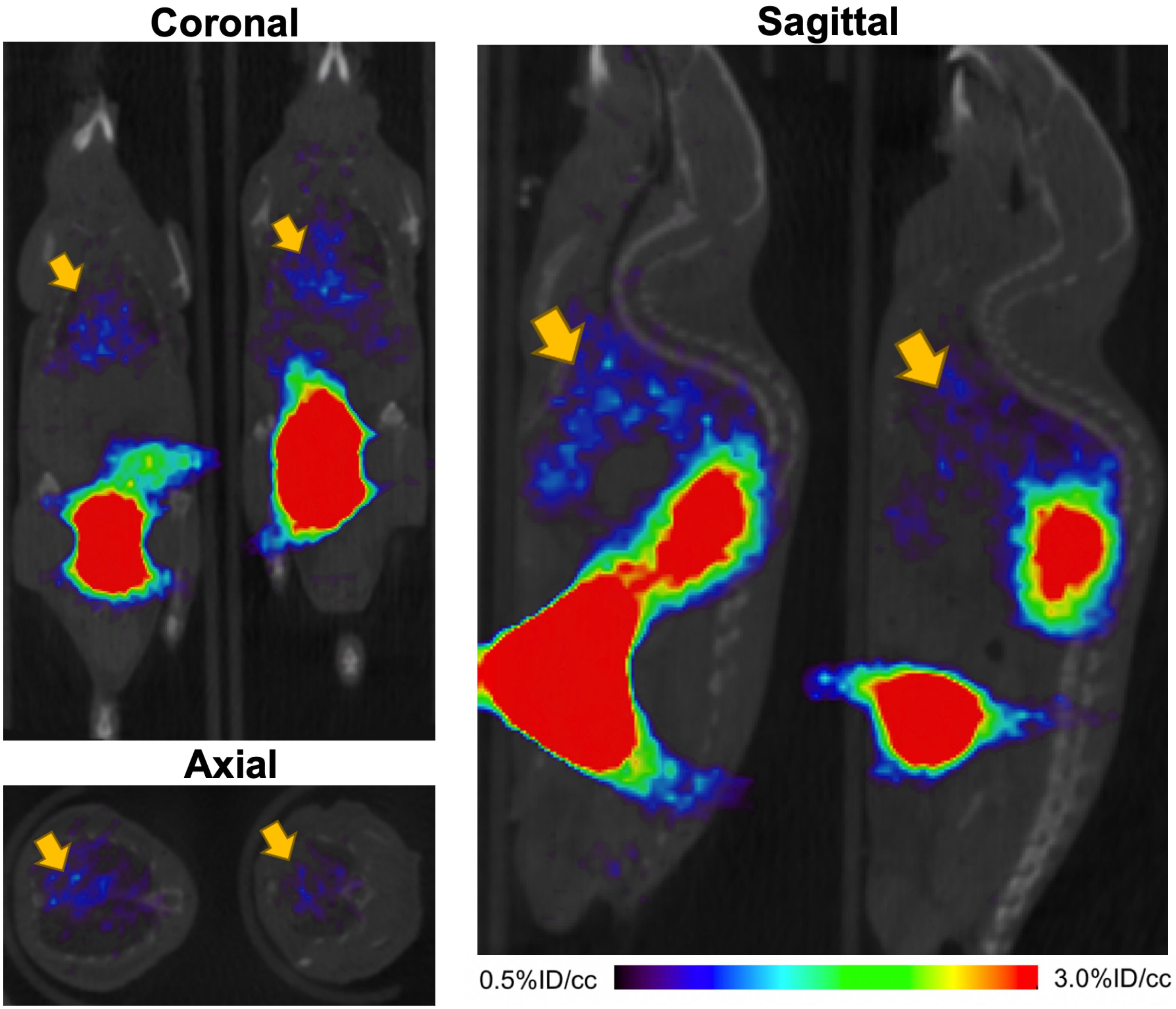

**Supp. Fig. 3:** Additional images of [$^{68}$Ga]-NOTA-ACE2pep4 in hACE2 transgenic mice, obtained via single time-point acquisition.

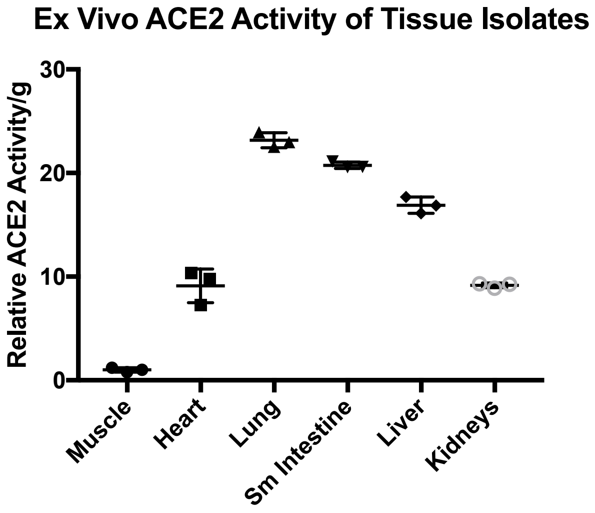

**Supp. Fig. 4:** Confirmation of ACE2 activity in tissue isolates, as correlation to reported *in vivo* data (Anaspec, Fremont CA).

Eurogentec

### ANASPEC PEPTIDE
#### QC Data Sheet

ANASPEC

For Research Use Only

| YOUR ORDER | SON | DATE |
|---|---|---|
| B001890494 | 1500055456 | 15-Jun-20 |

| CUSTOMER | ADDRESS / INSTITUTION |
|---|---|
| University of California San Francisco (UCSF) | UNITED STATES San Francisco |

| PEPTIDE NAME | LOT# | INTERIM | SCALE | # AMINO |
|---|---|---|---|---|
| 75386-1: (linear peptide) | **2056275** | 2071245 | Custom | 28 |

**SEQUENCE (N-Term → C-Term)**

**NOTA** - GGG DYS HCS PLR YYP WWK CTY PDP EGG G - **NH2**

| PHYSICOCHEMICAL PROPERTIES | | REGULAR AA PROPERTIES | | | | | |
|---|---|---|---|---|---|---|---|
| 1A 280 [mg/ml] * | 0.2 | Charged AA | D,H,R,K,E | 6 | Polar AA | D,S,H,R,K,T,E | 9 |
| Charged at pH 7 * | 0.0 | Acid AA | D,E | 3 | Hydrophobic AA | G,Y,C,P,L,W | 1 |
| Isoelectric Point * | 6.7 | Basic AA | H,R,K | 3 | | | |

*(* Theoretical values)*

**QC DATA**

| Attribute | Test method | Acceptance criteria | Result |
|---|---|---|---|
| Appearance | Visual | Report result | White Powder |
| % Peak Area by HPLC | HPLC | ≥ 90 % | 96 % |
| Identity | MS | 3433.9 ± 0.2 % | 3435.1 |

**DELIVERABLE**

**Format** Dried

**Aliquoting**
Number of Aliquots 1
Qty by Aliquot (mg) 10 mg

LOT# 2056275 10 mg
(linear peptide)
For Laboratory Use Only
ANASPEC

**DELIVERY CONDITION**
Room temperature

**STORAGE CONDITION**
-20 °C, dry

**COMMENTS**

#### PEPTIDE RECONSTITUTION AND STORAGE

Please read the entire section before proceeding with the solubilization of your custom peptide.

Peptides are shipped at ambient temperature as a lyophilized powder. Upon receipt store them at -20°C. Allow the vial to equilibrate to room temperature prior to opening.

Peptide solubility is highly dependent on the sequence. Peptides that are more hydrophobic (high propensity of A, F, G, V, L, I, M, W, P) in nature, will require an organic solvent in order to dissolve. Peptides that are acidic in nature (high propensity of D, E in the peptide sequence) require a basic aqueous buffer to dissolve, while peptides that are basic in nature (high propensity of K, H, and R) require an acidic aqueous buffer to dissolve.

To reconstitute a hydrophobic peptide, add 100 µL DMSO and sonicate until a homogenous solution forms. Next, add your buffer of choice to form a 1 mg/mL solution (a higher concentration of peptide will require a greater amount of DMSO). To reconstitute basic or acidic peptides, add 1 mL of the appropriate buffer to the peptide and sonicate to ensure a homogenous solution forms.

Reconstituted peptides can be stored frozen at -20°C for short period of time, but it is advisable to prepare multiple aliquots to avoid multiple freeze thaw cycles. We recommend that all aliquoted solutions be lyophilized if the peptide is going to be stored for extended periods of time at -20 °C.

Additionally, please note that peptides with a high propensity of basic residues (R, K, H) in their sequence may undergo a physical change from solid powder to an oil (via moisture absorption). This physical change does not affect the purity or functionality of the peptide.

Nomenclature used for the sequence termini:
- N-terminus: H means free amine ($NH_2$-), Ac mean acetyl [CH3C(O)-NH-], Pyr means pyroglutamic acid
- C-terminus: OH means free acid (-COOH), $NH_2$ means amide [-CONH$_2$]

Modifications on the side chain of amino acids are depicted in the parenthesis after the corresponding amino acid. For example; phosphorylated serine = S($PO_3H_2$) or epsilon-N-acetylated lysine = K(Ac)

#### TECHNICAL SUPPORT

If you have any questions feel free to call our Technical Support Centre

**EUROPE**
✆ 00 800 666 00 123 (European toll free number),
✉
Kaneka Eurogentec S.A. Liège Science Park
Rue Bois Saint-Jean 5 - 4102 SERAING BELGIUM
 Web: www.eurogentec.com
RPM Liège T.V.A.-(BE)-0427.348.346 - ING Belgique Bank - IBAN: BE86 3400 2118 6050 BIC: BBRUBEBB

**NORTH AMERICA**
✆ +1 800 452-5530 (American toll free number),
✉
AnaSpec, Inc. 34801 Campus Drive
Fremont, CA 94555 - USA
 Web: www.anaspec.com

### Chromatogram and Results

#### Injection Details

| | | | |
|---|---|---|---|
| *Injection Name:* | **2071245** | *Run Time (min):* | **12.00** |
| *Vial Number:* | **GA1** | *Injection Volume:* | **1.00** |
| *Injection Type:* | **Unknown** | *Channel:* | **UV_VIS_1** |
| *Column:* | **C18,100X4.6mm,H17-231434** | *Wavelength:* | **220** |
| *Instrument Method:* | **5-60%B-7minsExtended,Column(6-1)-0.7ml-30C** | *Bandwidth:* | **4** |
| *Processing Method:* | **test** | *Instrument No.* | **QC-HPLC-10** |
| *Injection Date/Time:* | **15/Jun/20 11:31** | *Sample Weight:* | |

#### Chromatogram

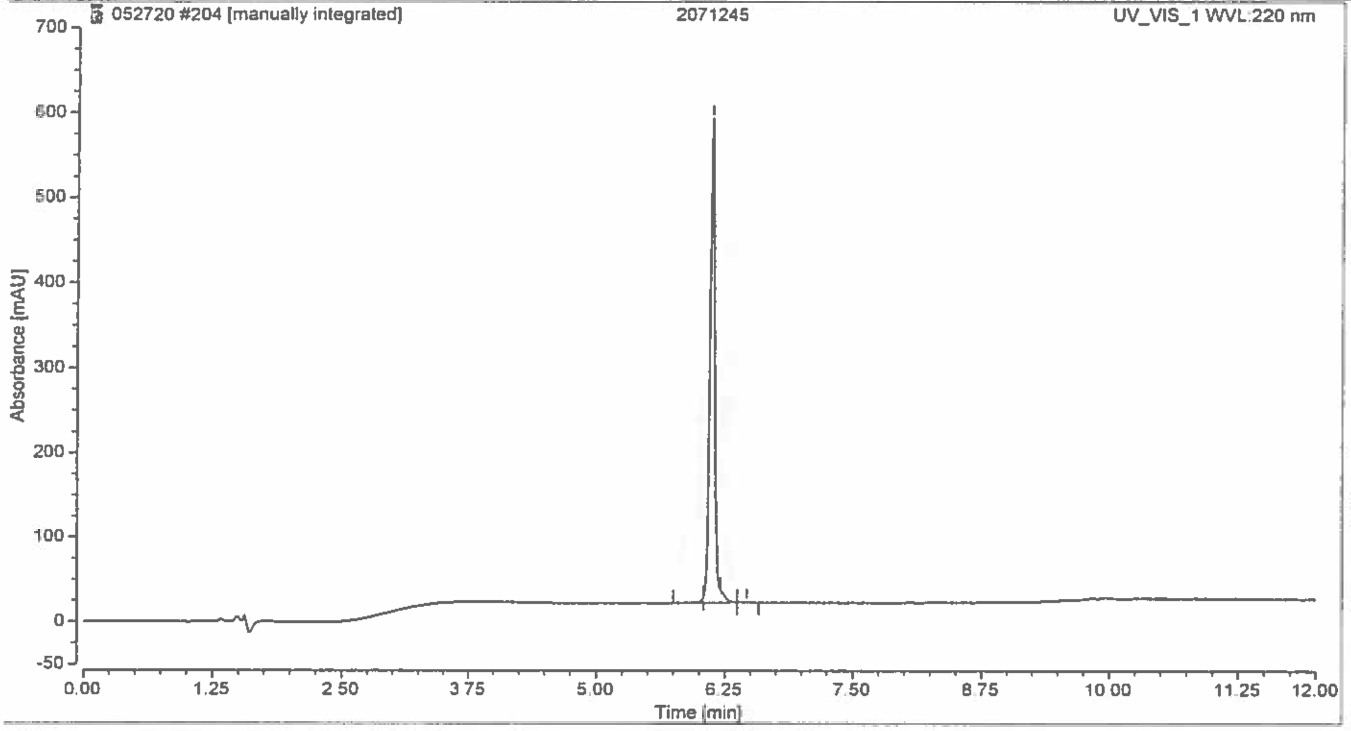

#### Integration Results

| No. | Retention Time<br>min | Area<br>mAU*min | Height<br>mAU | Relative Area<br>% |
|---|---|---|---|---|
| 1 | 6.040 | 0.204 | 6.008 | 0.64 |
| 2 | 6.120 | 30.586 | 571.320 | 96.41 |
| 3 | 6.207 | 0.807 | 15.001 | 2.55 |
| 4 | 6.463 | 0.128 | 1.142 | 0.40 |
| **Total:** | | **31.724** | **593.470** | **100.00** |

C:\Xcalibur\Data\2071245 06/12/20 12:30:23

06/12/20

2071245 #35-120 RT: 0.22-0.77 AV: 29 NL: 4.36E5
F: ITMS + c ESI E Full ms [50.00-2000.00]

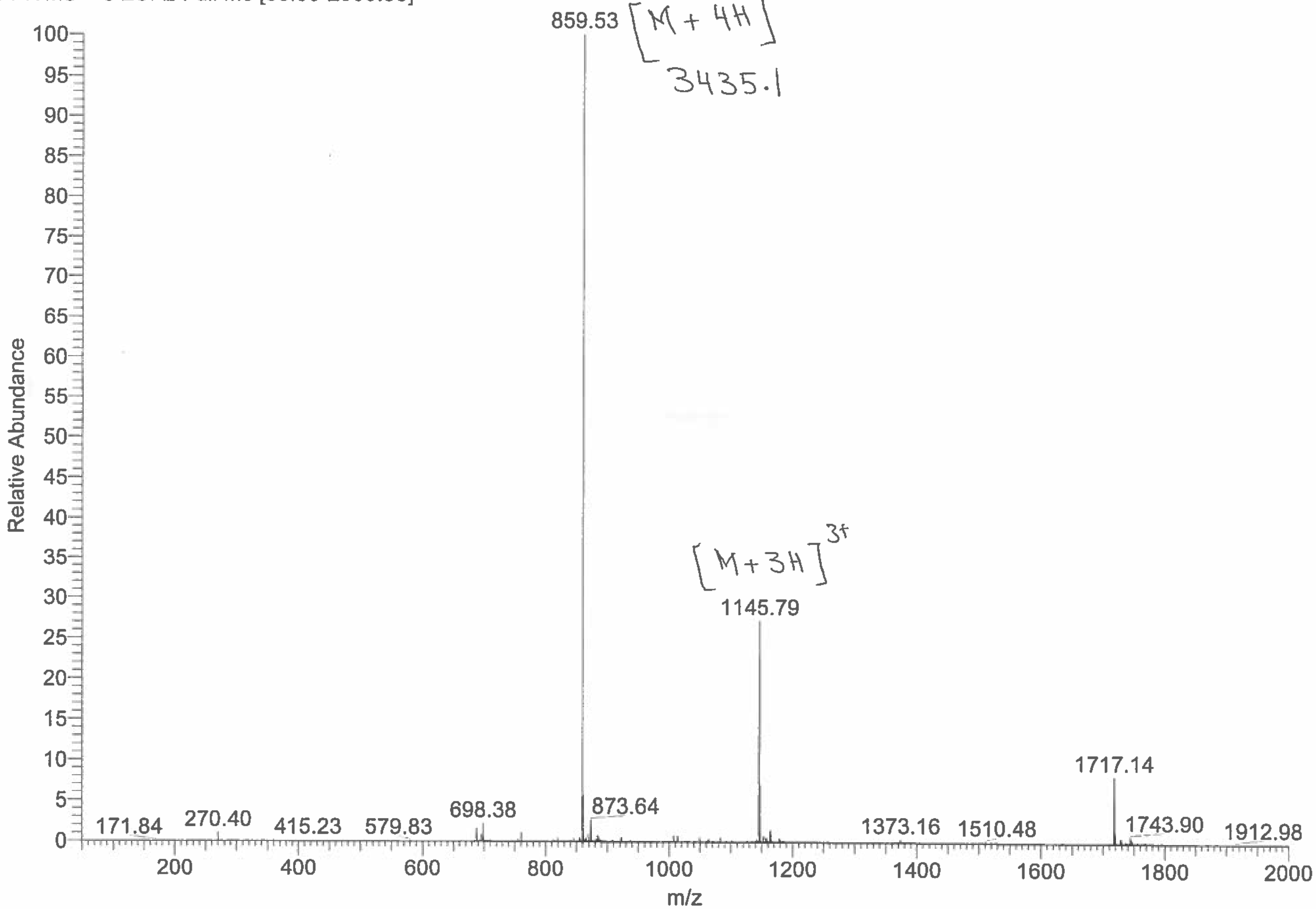

### ANASPEC PEPTIDE
#### QC Data Sheet

Eurogentec — ANASPEC — For Research Use Only

| YOUR ORDER | SON | DATE |
|---|---|---|
| B001890494 | 1500055456 | 16-Jun-20 |

| CUSTOMER | ADDRESS / INSTITUTION |
|---|---|
| University of California San Francisco (UCSF) | UNITED STATES San Francisco |

| PEPTIDE NAME | LOT# | INTERIM | SCALE | # AMINO |
|---|---|---|---|---|
| 75386-2: (disulfide bridge) | 2056276 | 2071246 | Custom | 28 |

**SEQUENCE (N-Term → C-Term)**

**NOTA** - GGG DYS HC(S-)S PLR YYP WWK C(S-)TY PDP EGG G - **NH2** , with disulfide bridge

| PHYSICOCHEMICAL PROPERTIES | |
|---|---|
| 1A 280 [mg/ml] * | 0.2 |
| Charged at pH 7 * | 0.0 |
| Isoelectric Point * | 6.7 |

(* Theoretical values)

| REGULAR AA PROPERTIES | | | | | |
|---|---|---|---|---|---|
| Charged AA | D,H,R,K,E | 6 | Polar AA | D,S,H,R,K,T,E | 9 |
| Acid AA | D,E | 3 | Hydrophobic AA | G,Y,P,L,W | 1 |
| Basic AA | H,R,K | 3 | | | |

**QC DATA**

| Attribute | Test method | Acceptance criteria | Result |
|---|---|---|---|
| Appearance | Visual | Report result | White Powder |
| % Peak Area by HPLC | HPLC | ≥ 95 % | 95 % |
| Identity | MS | 3431.9 ± 0.2 % | 3432.6 |

**DELIVERABLE**

**Format** Dried

**Aliquoting**
Number of Aliquots 1
Qty by Aliquot (mg) 10 mg

LOT: 2056276 10 mg
(disulfide bridge)
For Laboratory Use Only
ANASPEC

**DELIVERY CONDITION**
Room temperature

**STORAGE CONDITION**
-20 °C, dry

**COMMENTS**

#### PEPTIDE RECONSTITUTION AND STORAGE

Please read the entire section before proceeding with the solubilization of your custom peptide.

Peptides are shipped at ambient temperature as a lyophilized powder. Upon receipt store them at -20°C. Allow the vial to equilibrate to room temperature prior to opening.

Peptide solubility is highly dependent on the sequence. Peptides that are more hydrophobic (high propensity of A, F, G, V, L, I, M, W, P) in nature, will require an organic solvent in order to dissolve. Peptides that are acidic in nature (high propensity of D, E in the peptide sequence) require a basic aqueous buffer to dissolve, while peptides that are basic in nature (high propensity of K, H, and R) require an acidic aqueous buffer to dissolve.

To reconstitute a hydrophobic peptide, add 100 µL DMSO and sonicate until a homogenous solution forms. Next, add your buffer of choice to form a 1 mg/mL solution (a higher concentration of peptide will require a greater amount of DMSO). To reconstitute basic or acidic peptides, add 1 mL of the appropriate buffer to the peptide and sonicate to ensure a homogenous solution forms.

Reconstituted peptides can be stored frozen at -20°C for short period of time, but it is advisable to prepare multiple aliquots to avoid multiple freeze thaw cycles. We recommend that all aliquoted solutions be lyophilized if the peptide is going to be stored for extended periods of time at -20 °C.

Additionally, please note that peptides with a high propensity of basic residues (R, K, H) in their sequence may undergo a physical change from solid powder to an oil (via moisture absorption). This physical change does not affect the purity or functionality of the peptide.

Nomenclature used for the sequence termini:

N-terminus: H means free amine ($NH_2$-), Ac mean acetyl [CH3C(O)-NH-], Pyr means pyroglutamic acid
C-terminus: OH means free acid (-COOH), $NH_2$ means amide [-$CONH_2$]

Modifications on the side chain of amino acids are depicted in the parenthesis after the corresponding amino acid. For example; phosphorylated serine = S($PO_3H_2$) or epsilon-N-acetylated lysine = K(Ac)

#### TECHNICAL SUPPORT

If you have any questions feel free to call our Technical Support Centre

**EUROPE**

✆ 00 800 666 00 123 (European toll free number),
✉

Kaneka Eurogentec S.A. Liège Science Park
Rue Bois Saint-Jean 5 - 4102 SERAING BELGIUM
 Web: www.eurogentec.com
RPM Liège T.V.A.-(BE)-0427.348.346 - ING Belgique Bank - IBAN: BE86 3400 2118 6050 BIC: BBRUBEBB

**NORTH AMERICA**

✆ +1 800 452-5530 (American toll free number),
✉

AnaSpec, Inc. 34801 Campus Drive
Fremont, CA 94555 - USA
 Web www.anaspec.com

### Chromatogram and Results

#### Injection Details

| | | | |
|---|---|---|---|
| *Injection Name:* | **2071246** | *Run Time (min):* | **12.00** |
| *Vial Number:* | **GC2** | *Injection Volume:* | **1.00** |
| *Injection Type:* | **Unknown** | *Channel:* | **UV_VIS_1** |
| *Column:* | **C18,100X4.6mm,H17-231434** | *Wavelength:* | **220** |
| *Instrument Method:* | **5-60%B-7minsExtended,Column(6-1)-0.7ml-30C** | *Bandwidth:* | **4** |
| *Processing Method:* | **test** | *Instrument No.* | **QC-HPLC-10** |
| *Injection Date/Time:* | **16/Jun/20 13:47** | *Sample Weight:* | |

#### Chromatogram

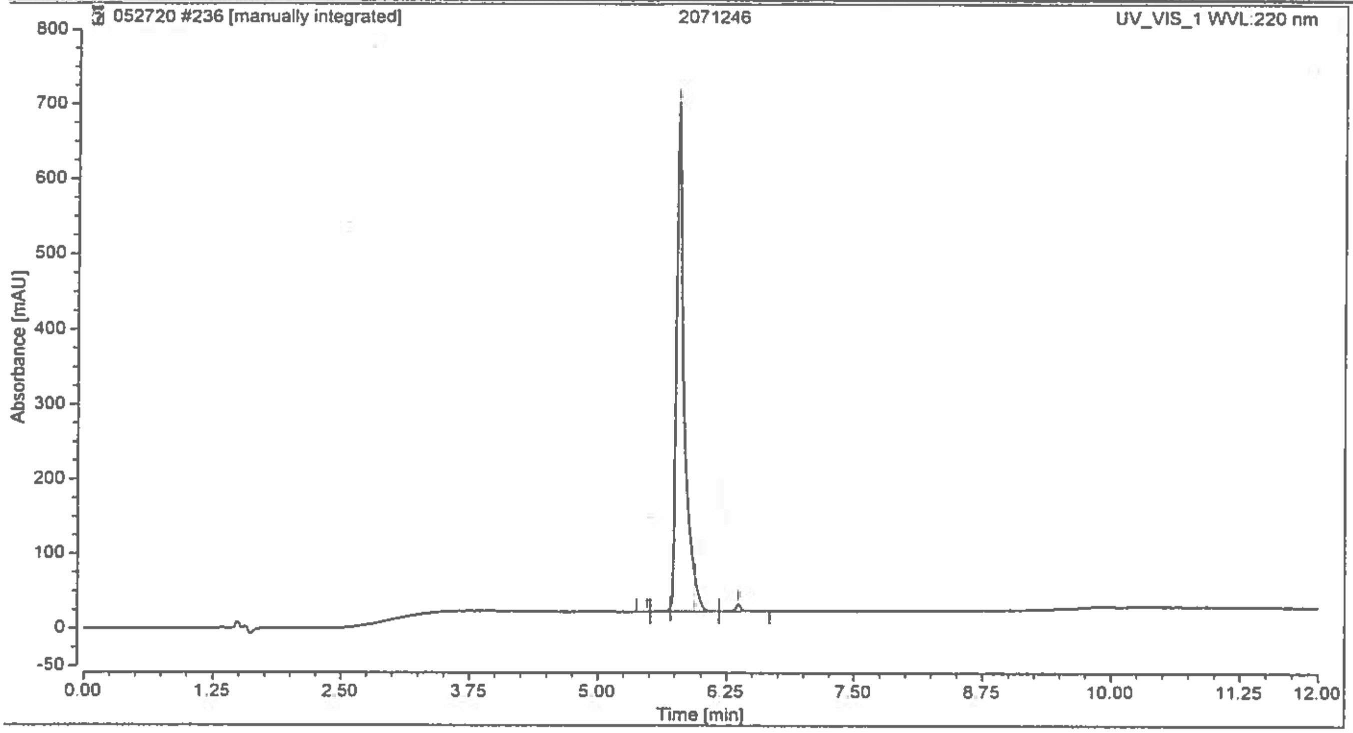

#### Integration Results

| No. | Retention Time min | Area mAU*min | Height mAU | Relative Area % |
|---|---|---|---|---|
| 1 | 5.477 | 0.003 | 0.104 | 0.01 |
| 2 | 5.707 | 0.096 | 3.887 | 0.17 |
| 3 | 5.773 | 55.557 | 679.939 | 95.33 |
| 4 | 5.937 | 2.010 | 47.591 | 3.45 |
| 5 | 6.367 | 0.611 | 10.131 | 1.05 |
| **Total:** | | **58.277** | **741.651** | **100.00** |

C:\Xcalibur\Data\2071246 06/16/20 11:13:43

06/16/2020

2071246 #35-114 RT: 0.22-0.73 AV: 27 NL: 2.03E5
F: ITMS + c ESI E Full ms [50.00-2000.00]

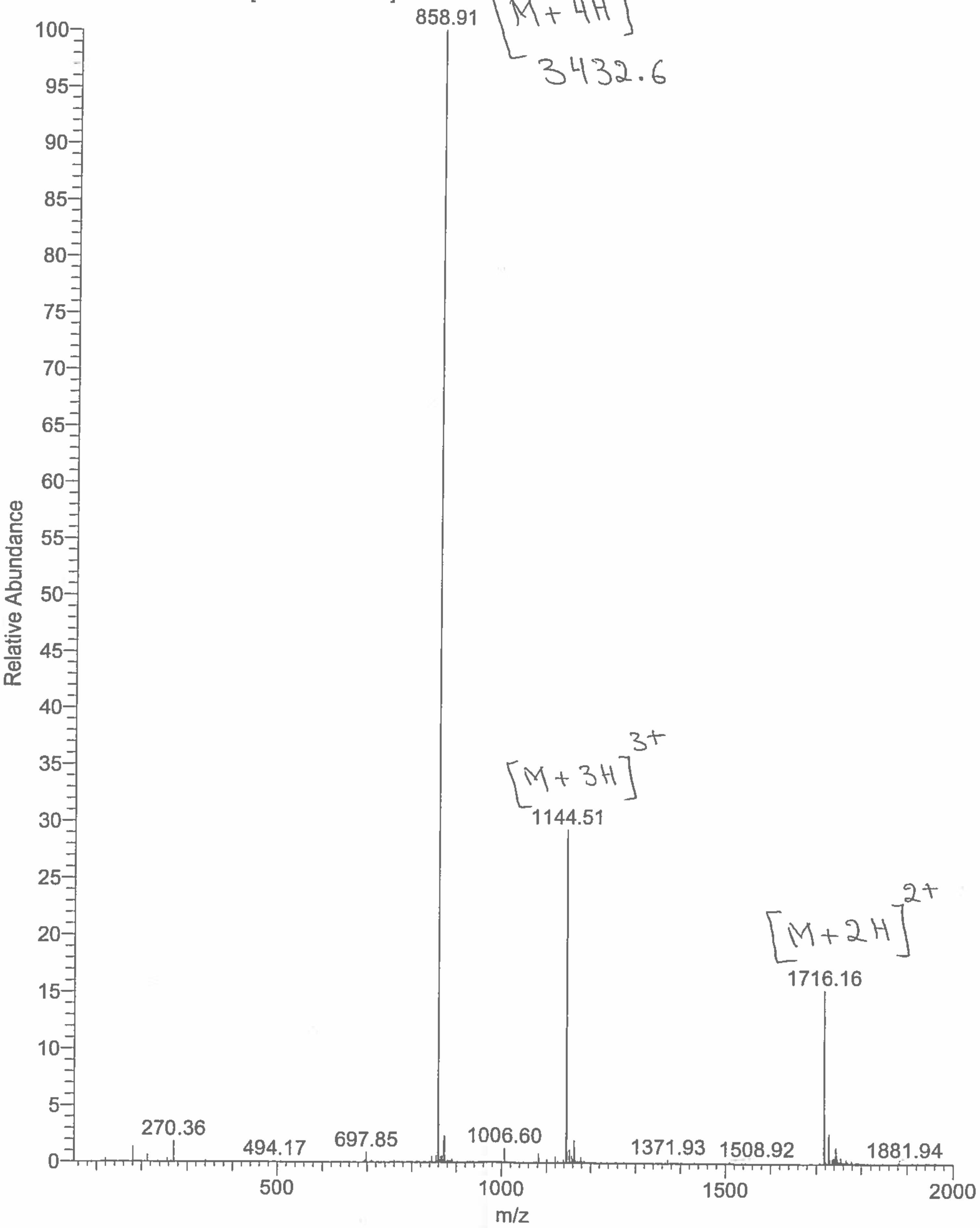

Eurogentec

### ANASPEC PEPTIDE
#### QC Data Sheet

ANASPEC

For Research Use Only

| YOUR ORDER | SON | DATE |
|---|---|---|
| B001890494 | 1500055456 | 10-Jun-20 |

| CUSTOMER | ADDRESS / INSTITUTION |
|---|---|
| University of California San Francisco (UCSF) | UNITED STATES San Francisco |

| PEPTIDE NAME | LOT# | INTERIM | SCALE | # AMINO |
|---|---|---|---|---|
| 75386-3: (linear peptide) | **2056277** | 2071247 | Custom | 26 |

**SEQUENCE (N-Term → C-Term)**

**NOTA** -Ahx- DY SHC SPL RYY PWW KCT YPD PEG GG - **NH2**
Ahx=6-aminohexanoic acid linker

| PHYSICOCHEMICAL PROPERTIES | |
|---|---|
| 1A 280 [mg/ml] * | 0.2 |
| Charged at pH 7 * | 0.0 |
| Isoelectric Point * | 6.7 |

(* Theoretical values)

| REGULAR AA PROPERTIES | | | | | |
|---|---|---|---|---|---|
| Charged AA | D,H,R,K,E | 6 | Polar AA | D,S,H,R,K,T,E | 9 |
| Acid AA | D,E | 3 | Hydrophobic AA | Y,C,P,L,W,G,-Ahx- | 1 |
| Basic AA | H,R,K,-Ahx- | 4 | | | |

**QC DATA**

| Attribute | Test method | Acceptance criteria | Result |
|---|---|---|---|
| Appearance | Visual | Report result | White Powder |
| % Peak Area by HPLC | HPLC | ≥ 90 % | 96 % |
| Identity | MS | 3375.9 ± 0.2 % | 3376.8 |

**DELIVERABLE**

**Format** Dried

**Aliquoting**
Number of Aliquots 1
Qty by Aliquot (mg) 10 mg

LOT# 2056277 10 mg
(linear peptide)
For Laboratory Use Only
ANASPEC

**DELIVERY CONDITION**
Room temperature

**STORAGE CONDITION**
-20 °C, dry

**COMMENTS**

**PEPTIDE RECONSTITUTION AND STORAGE**

Please read the entire section before proceeding with the solubilization of your custom peptide.

Peptides are shipped at ambient temperature as a lyophilized powder. Upon receipt store them at -20°C. Allow the vial to equilibrate to room temperature prior to opening.

Peptide solubility is highly dependent on the sequence. Peptides that are more hydrophobic (high propensity of A, F, G, V, L, I, M, W, P) in nature, will require an organic solvent in order to dissolve. Peptides that are acidic in nature (high propensity of D, E in the peptide sequence) require a basic aqueous buffer to dissolve, while peptides that are basic in nature (high propensity of K, H, and R) require an acidic aqueous buffer to dissolve.

To reconstitute a hydrophobic peptide, add 100 µL DMSO and sonicate until a homogenous solution forms. Next, add your buffer of choice to form a 1 mg/mL solution (a higher concentration of peptide will require a greater amount of DMSO). To reconstitute basic or acidic peptides, add 1 mL of the appropriate buffer to the peptide and sonicate to ensure a homogenous solution forms.

Reconstituted peptides can be stored frozen at -20°C for short period of time, but it is advisable to prepare multiple aliquots to avoid multiple freeze thaw cycles. We recommend that all aliquoted solutions be lyophilized if the peptide is going to be stored for extended periods of time at -20 °C.

Additionally, please note that peptides with a high propensity of basic residues (R, K, H) in their sequence may undergo a physical change from solid powder to an oil (via moisture absorption). This physical change does not affect the purity or functionality of the peptide.

Nomenclature used for the sequence termini:
N-terminus: H means free amine ($NH_2$-), Ac mean acetyl [CH3C(O)-NH-], Pyr means pyroglutamic acid
C-terminus: OH means free acid (-COOH), $NH_2$ means amide [-$CONH_2$]

Modifications on the side chain of amino acids are depicted in the parenthesis after the corresponding amino acid. For example; phosphorylated serine = S($PO_3H_2$) or epsilon-N-acetylated lysine = K(Ac)

**TECHNICAL SUPPORT**

If you have any questions feel free to call our Technical Support Centre

**EUROPE**

✆ 00 800 666 00 123 (European toll free number),
✉

Kaneka Eurogentec S.A. Liège Science Park
Rue Bois Saint-Jean 5 - 4102 SERAING BELGIUM
 Web: www.eurogentec.com
RPM Liège T.V.A.-(BE)-0427.348.346 - ING Belgique Bank - IBAN: BE86 3400 2118 6050 BIC: BBRUBEBB

**NORTH AMERICA**

✆ +1 800 452-5530 (American toll free number),
✉

AnaSpec, Inc. 34801 Campus Drive
Fremont, CA 94555 - USA
 Web: www.anaspec.com

### Chromatogram and Results

#### Injection Details

| | | | |
|---|---|---|---|
| *Injection Name:* | **2071247** | *Run Time (min):* | **12.00** |
| *Vial Number:* | **BA1** | *Injection Volume:* | **1.50** |
| *Injection Type:* | **Unknown** | *Channel:* | **UV_VIS_1** |
| *Column:* | **C18,100X4.6mm,H17-231434** | *Wavelength:* | **220** |
| *Instrument Method:* | **5-60%B-7minsExtended,Column(6-1)-0.7ml-30C** | *Bandwidth:* | **4** |
| *Processing Method:* | **test** | *Instrument No.* | **QC-HPLC-10** |
| *Injection Date/Time:* | **10/Jun/20 13:56** | *Sample Weight:* | |

#### Chromatogram

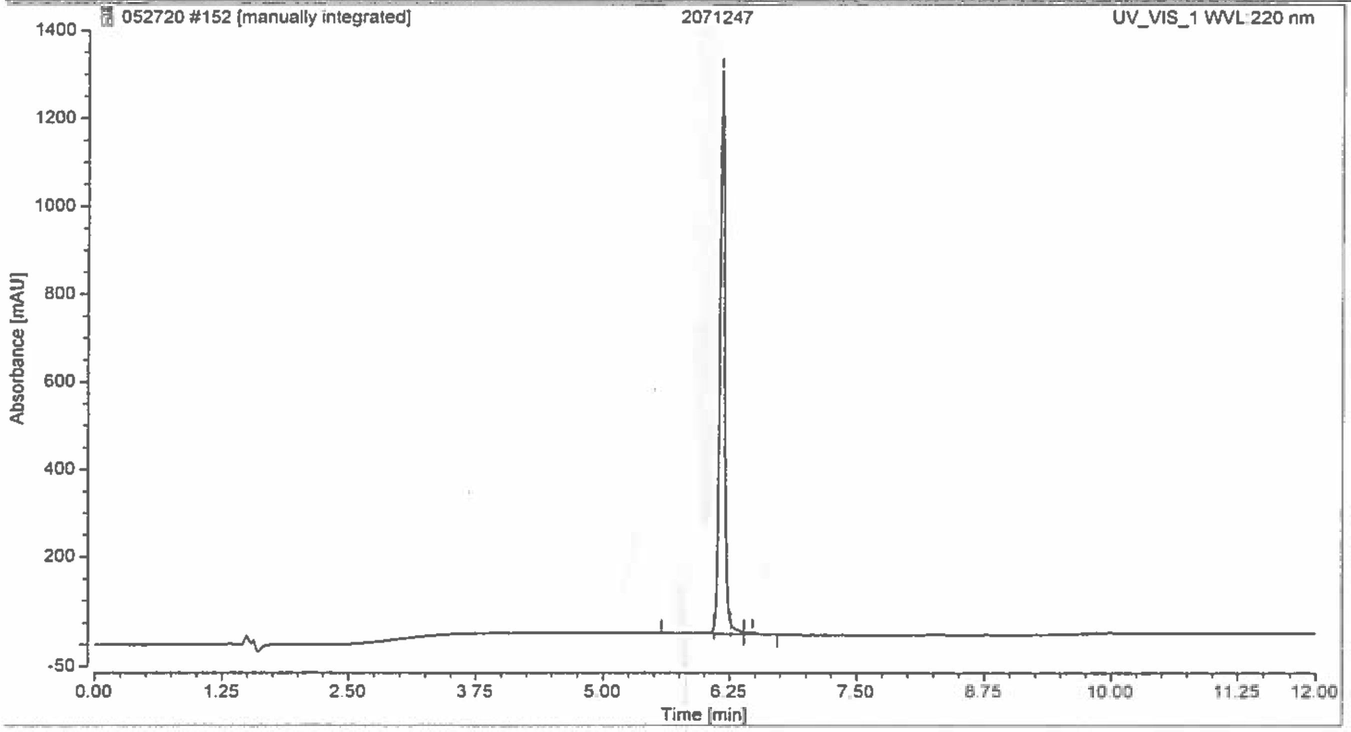

#### Integration Results

| No. | Retention Time min | Area mAU*min | Height mAU | Relative Area % |
|---|---|---|---|---|
| 1 | 6.097 | 0.652 | 16.476 | 0.91 |
| 2 | 6.170 | 69.280 | 1283.037 | 96.41 |
| 3 | 6.260 | 1.376 | 21.965 | 1.92 |
| 4 | 6.473 | 0.555 | 5.212 | 0.77 |
| **Total:** | | **71.863** | **1326.691** | **100.00** |

C:\Xcalibur\Data\2071247 06/10/20 13:53:34

06/10/20

2071247 #35-109 RT: 0.22-0.69 AV: 25 NL: 2.70E5
F: ITMS + c ESI E Full ms [50.00-2000.00]

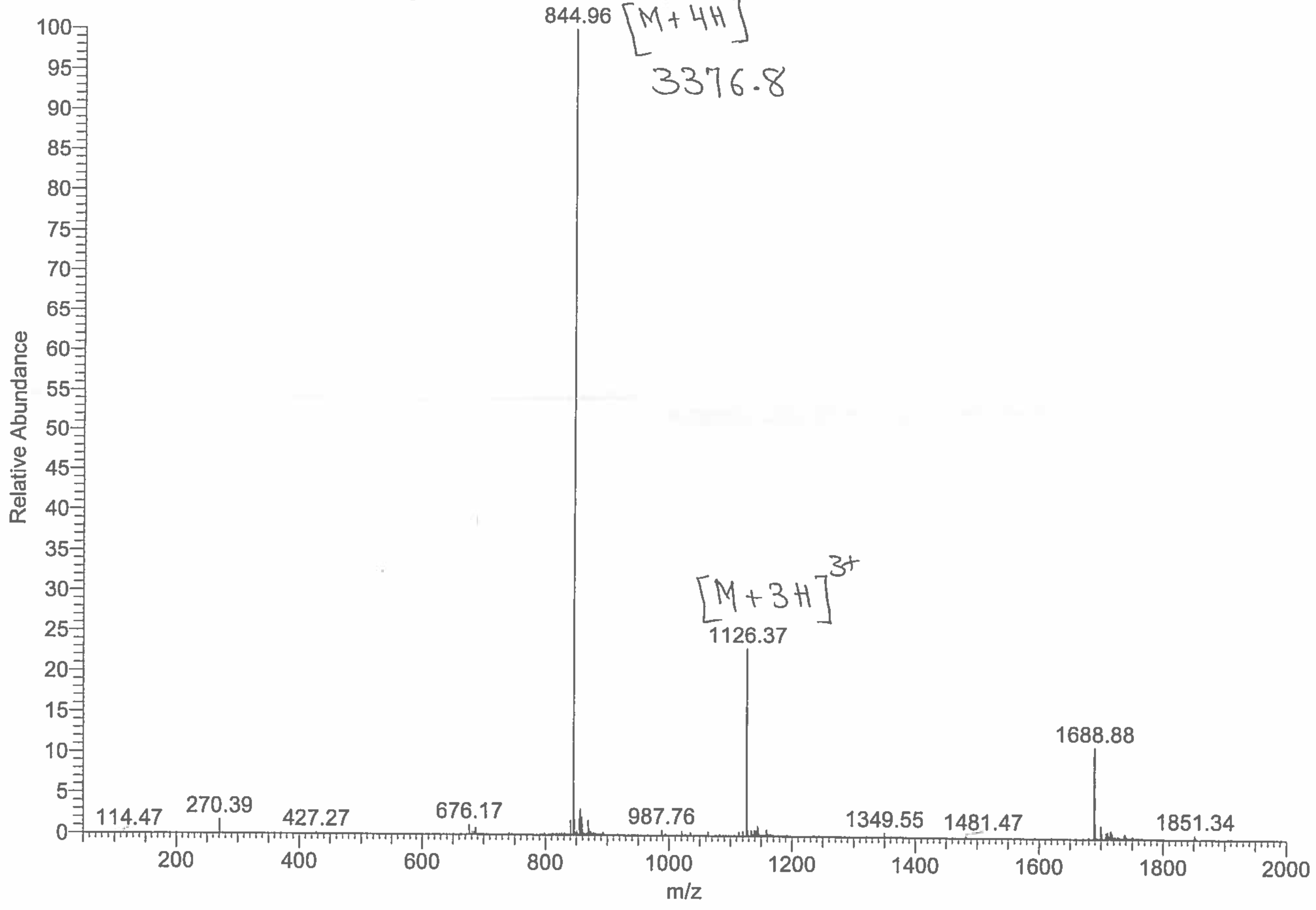

Eurogentec

### ANASPEC PEPTIDE
#### QC Data Sheet

ANASPEC

For Research Use Only

| YOUR ORDER | SON | DATE |
|---|---|---|
| B001890494 | 1500055456 | 10-Jun-20 |

| CUSTOMER | ADDRESS / INSTITUTION |
|---|---|
| University of California San Francisco (UCSF) | UNITED STATES San Francisco |

| PEPTIDE NAME | LOT# | INTERIM | SCALE | # AMINO |
|---|---|---|---|---|
| 75386-4: (disulfide bridge) | **2056278** | 2071248 | Custom | 26 |

**SEQUENCE (N-Term → C-Term)**

**NOTA-** Ahx- DY SHC(S-) SPL RYY PWW KC(S-)T YPD PEG GG **- NH2** , with disulfide bridge
Ahx=6-aminohexanoic acid linker

| PHYSICOCHEMICAL PROPERTIES | |
|---|---|
| 1A 280 [mg/ml] * | 0.2 |
| Charged at pH 7 * | 0.0 |
| Isoelectric Point * | 6.7 |

(* Theoretical values)

| REGULAR AA PROPERTIES | | | | | |
|---|---|---|---|---|---|
| Charged AA | D,H,R,K,E | 6 | Polar AA | D,S,H,R,K,T,E | 9 |
| Acid AA | D,E | 3 | Hydrophobic AA | Y,P,L,W,G,-Ahx- | 1 |
| Basic AA | H,R,K,-Ahx- | 4 | | | |

**QC DATA**

| Attribute | Test method | Acceptance criteria | Result |
|---|---|---|---|
| Appearance | Visual | Report result | White Powder |
| % Peak Area by HPLC | HPLC | ≥ 95 % | 95 % |
| Identity | MS | 3373.8 ± 0.2 % | 3374.8 |

**DELIVERABLE**

**Format** Dried

**Aliquoting**
Number of Aliquots 1
Qty by Aliquot (mg) 10 mg

**DELIVERY CONDITION**
Room temperature

**STORAGE CONDITION**
-20 °C, dry

**COMMENTS**

LOT# 2056278 10 mg
(Disulfide bridge)
For Laboratory Use Only
ANASPEC

**PEPTIDE RECONSTITUTION AND STORAGE**

Please read the entire section before proceeding with the solubilization of your custom peptide.

Peptides are shipped at ambient temperature as a lyophilized powder. Upon receipt store them at -20°C. Allow the vial to equilibrate to room temperature prior to opening.

Peptide solubility is highly dependent on the sequence. Peptides that are more hydrophobic (high propensity of A, F, G, V, L, I, M, W, P) in nature, will require an organic solvent in order to dissolve. Peptides that are acidic in nature (high propensity of D, E in the peptide sequence) require a basic aqueous buffer to dissolve, while peptides that are basic in nature (high propensity of K, H, and R) require an acidic aqueous buffer to dissolve.

To reconstitute a hydrophobic peptide, add 100 µL DMSO and sonicate until a homogenous solution forms. Next, add your buffer of choice to form a 1 mg/mL solution (a higher concentration of peptide will require a greater amount of DMSO). To reconstitute basic or acidic peptides, add 1 mL of the appropriate buffer to the peptide and sonicate to ensure a homogenous solution forms.

Reconstituted peptides can be stored frozen at -20°C for short period of time, but it is advisable to prepare multiple aliquots to avoid multiple freeze thaw cycles. We recommend that all aliquoted solutions be lyophilized if the peptide is going to be stored for extended periods of time at -20 °C.

Additionally, please note that peptides with a high propensity of basic residues (R, K, H) in their sequence may undergo a physical change from solid powder to an oil (via moisture absorption). This physical change does not affect the purity or functionality of the peptide.

Nomenclature used for the sequence termini:
N-terminus: H means free amine ($NH_2$-), Ac mean acetyl [$CH3C(O)$-NH-], Pyr means pyroglutamic acid
C-terminus: OH means free acid (-COOH), $NH_2$ means amide [-$CONH_2$]

Modifications on the side chain of amino acids are depicted in the parenthesis after the corresponding amino acid. For example; phosphorylated serine = S($PO_3H_2$) or epsilon-N-acetylated lysine = K(Ac)

**TECHNICAL SUPPORT**

If you have any questions feel free to call our Technical Support Centre

**EUROPE**
✆ 00 800 666 00 123 (European toll free number),
✉
Kaneka Eurogentec S.A. Liège Science Park
Rue Bois Saint-Jean 5 - 4102 SERAING BELGIUM
 Web: www.eurogentec.com
RPM Liège T.V.A.-(BE)-0427.348.346 - ING Belgique Bank - IBAN: BE86 3400 2118 6050 BIC: BBRUBEBB

**NORTH AMERICA**
✆ +1 800 452-5530 (American toll free number),
✉
AnaSpec, Inc. 34801 Campus Drive
Fremont, CA 94555 - USA
 Web: www.anaspec.com

### Chromatogram and Results

#### Injection Details

| | | | |
|---|---|---|---|
| *Injection Name:* | **2071248** | *Run Time (min):* | **12.00** |
| *Vial Number:* | **BA2** | *Injection Volume:* | **2.50** |
| *Injection Type:* | **Unknown** | *Channel:* | **UV_VIS_1** |
| *Column:* | **C18,100X4.6mm,H17-231434** | *Wavelength:* | **220** |
| *Instrument Method:* | **5-60%B-7minsExtended,Column(6-1)-0.7ml-30C** | *Bandwidth:* | **4** |
| *Processing Method:* | **test** | *Instrument No.* | **QC-HPLC-10** |
| *Injection Date/Time:* | **10/Jun/20 15:31** | *Sample Weight:* | |

#### Chromatogram

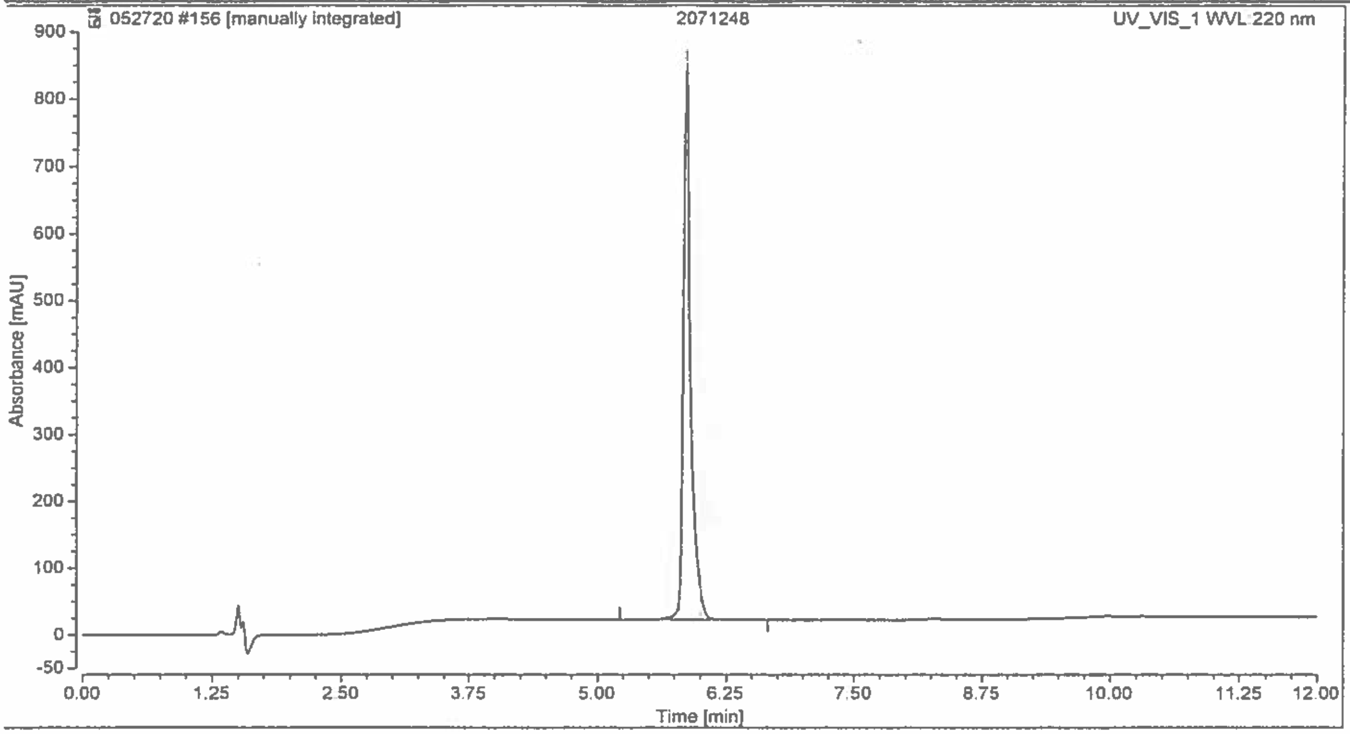

#### Integration Results

| No. | Retention Time min | Area mAU*min | Height mAU | Relative Area % |
|---|---|---|---|---|
| 1 | 5.803 | 1.364 | 49.670 | 2.05 |
| 2 | 5.853 | 63.551 | 829.478 | 95.29 |
| 3 | 5.997 | 1.779 | 47.305 | 2.67 |
| **Total:** | | **66.694** | **926.453** | **100.00** |

C:\Xcalibur\Data\2071248 06/10/20 13:56:59

2071248 #37-107 RT: 0.24-0.69 AV: 24 NL: 1.54E5
F: ITMS + c ESI E Full ms [50.00-2000.00]

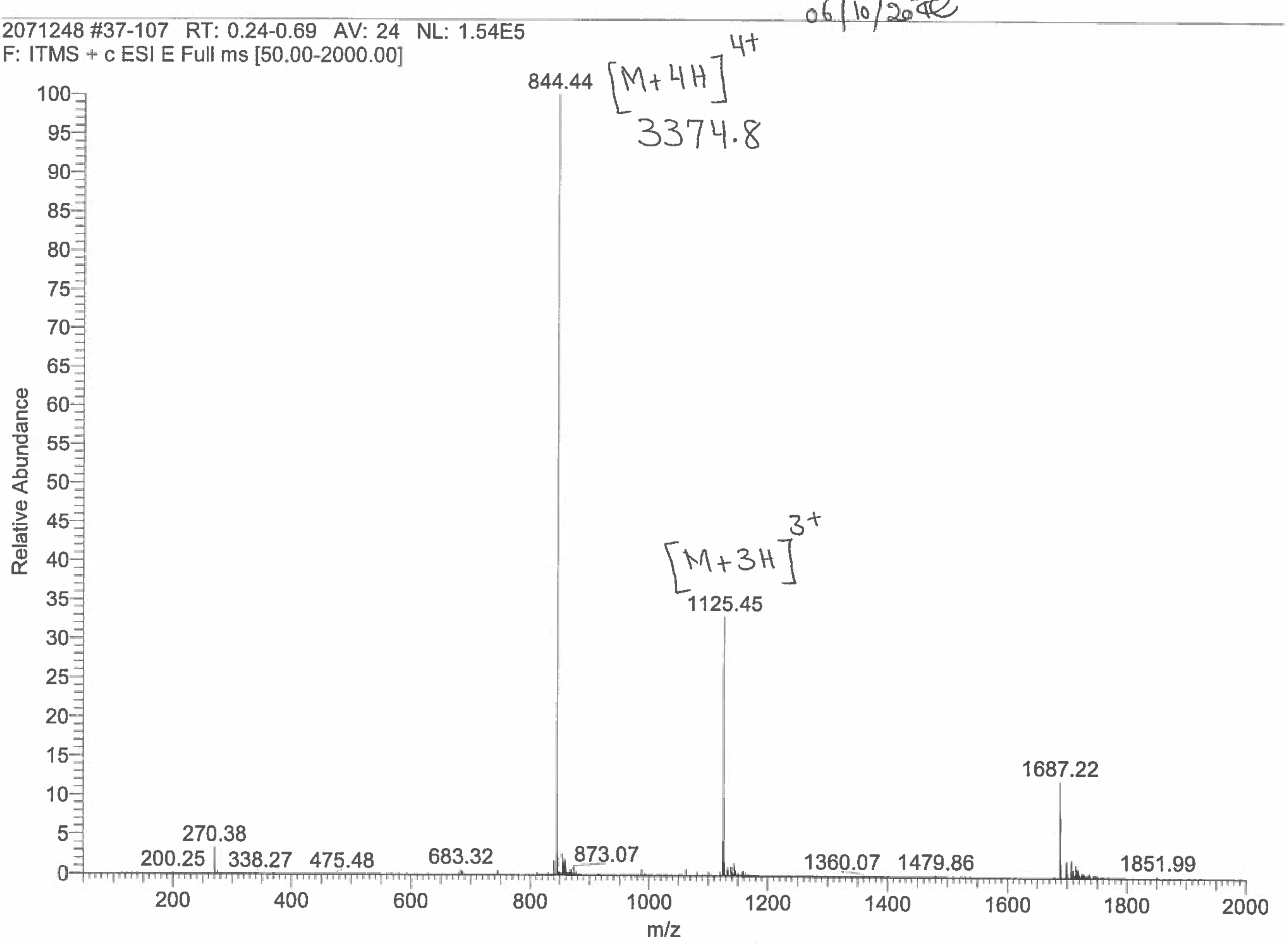

Eurogentec

### ANASPEC PEPTIDE
#### QC Data Sheet

ANASPEC

For Research Use Only

| YOUR ORDER | SON | DATE |
|---|---|---|
| B001890494 | 1500055456 | 19-Jun-20 |

| CUSTOMER | ADDRESS / INSTITUTION |
|---|---|
| University of California San Francisco (UCSF) | UNITED STATES San Francisco |

| PEPTIDE NAME | LOT# | INTERIM | SCALE | # AMINO |
|---|---|---|---|---|
| 75386-5: (linear peptide) | **2056279** | 2071249 | Custom | 26 |

**SEQUENCE (N-Term → C-Term)**

**NOTA -X-** DY SHC SPL RYY PWW KCT YPD PEG GG **- NH2**
X=AEEEA

| PHYSICOCHEMICAL PROPERTIES | | REGULAR AA PROPERTIES | | | | | |
|---|---|---|---|---|---|---|---|
| 1A 280 [mg/ml] * | 0.2 | Charged AA | D,H,R,K,E | 6 | Polar AA | D,S,H,R,K,T,E | 9 |
| Charged at pH 7 * | 0.0 | Acid AA | D,E | 3 | Hydrophobic AA | Y,C,P,L,W,G | 1 |
| Isoelectric Point * | 6.7 | Basic AA | H,R,K | 3 | | | |

(* Theoretical values)

**QC DATA**

| Attribute | Test method | Acceptance criteria | Result |
|---|---|---|---|
| Appearance | Visual | Report result | White Powder |
| % Peak Area by HPLC | HPLC | ≥ 90 % | 95 % |
| Identity | MS | 3451.9 ± 0.2 % | 3452.9 |

**DELIVERABLE**

**Format** Dried

**Aliquoting**
Number of Aliquots 1
Qty by Aliquot (mg) 10 mg

**DELIVERY CONDITION**
Room temperature

**STORAGE CONDITION**
-20 °C, dry

Lot# 2056279 10 mg
(linear peptide)
For Laboratory Use Only
ANASPEC

**COMMENTS**

#### PEPTIDE RECONSTITUTION AND STORAGE

Please read the entire section before proceeding with the solubilization of your custom peptide.

Peptides are shipped at ambient temperature as a lyophilized powder. Upon receipt store them at -20°C. Allow the vial to equilibrate to room temperature prior to opening.

Peptide solubility is highly dependent on the sequence. Peptides that are more hydrophobic (high propensity of A, F, G, V, L, I, M, W, P) in nature, will require an organic solvent in order to dissolve. Peptides that are acidic in nature (high propensity of D, E in the peptide sequence) require a basic aqueous buffer to dissolve, while peptides that are basic in nature (high propensity of K, H, and R) require an acidic aqueous buffer to dissolve.

To reconstitute a hydrophobic peptide, add 100 µL DMSO and sonicate until a homogenous solution forms. Next, add your buffer of choice to form a 1 mg/mL solution (a higher concentration of peptide will require a greater amount of DMSO). To reconstitute basic or acidic peptides, add 1 mL of the appropriate buffer to the peptide and sonicate to ensure a homogenous solution forms.

Reconstituted peptides can be stored frozen at -20°C for short period of time, but it is advisable to prepare multiple aliquots to avoid multiple freeze thaw cycles. We recommend that all aliquoted solutions be lyophilized if the peptide is going to be stored for extended periods of time at -20 °C.

Additionally, please note that peptides with a high propensity of basic residues (R, K, H) in their sequence may undergo a physical change from solid powder to an oil (via moisture absorption). This physical change does not affect the purity or functionality of the peptide.

Nomenclature used for the sequence termini:
N-terminus: H means free amine ($NH_2$-), Ac mean acetyl [CH3C(O)-NH-], Pyr means pyroglutamic acid
C-terminus: OH means free acid (-COOH), $NH_2$ means amide [-$CONH_2$]

Modifications on the side chain of amino acids are depicted in the parenthesis after the corresponding amino acid. For example; phosphorylated serine = S($PO_3H_2$) or epsilon-N-acetylated lysine = K(Ac)

#### TECHNICAL SUPPORT

If you have any questions feel free to call our Technical Support Centre

**EUROPE**
00 800 666 00 123 (European toll free number),

Kaneka Eurogentec S.A. Liège Science Park
Rue Bois Saint-Jean 5 - 4102 SERAING BELGIUM
 Web: www.eurogentec.com
RPM Liège T.V.A.-(BE)-0427.348 346 - ING Belgique Bank - IBAN: BE86 3400 2118 6050 BIC: BBRUBEBB

**NORTH AMERICA**
+1 800 452-5530 (American toll free number),

AnaSpec, Inc. 34801 Campus Drive
Fremont, CA 94555 - USA
 Web: www.anaspec.com

### Chromatogram and Results

#### Injection Details

| | | | |
|---|---|---|---|
| *Injection Name:* | **2071249** | *Run Time (min):* | **12.00** |
| *Vial Number:* | **GA3** | *Injection Volume:* | **1.00** |
| *Injection Type:* | **Unknown** | *Channel:* | **UV_VIS_1** |
| *Column:* | **C18,100X4.6mm,H17-231434** | *Wavelength:* | **220** |
| *Instrument Method:* | **5-60%B-7minsExtended,Column(6-1)-0.7ml-30C** | *Bandwidth:* | **4** |
| *Processing Method:* | **test** | *Instrument No.* | **QC-HPLC-10** |
| *Injection Date/Time:* | **18/Jun/20 16:03** | *Sample Weight:* | |

#### Chromatogram

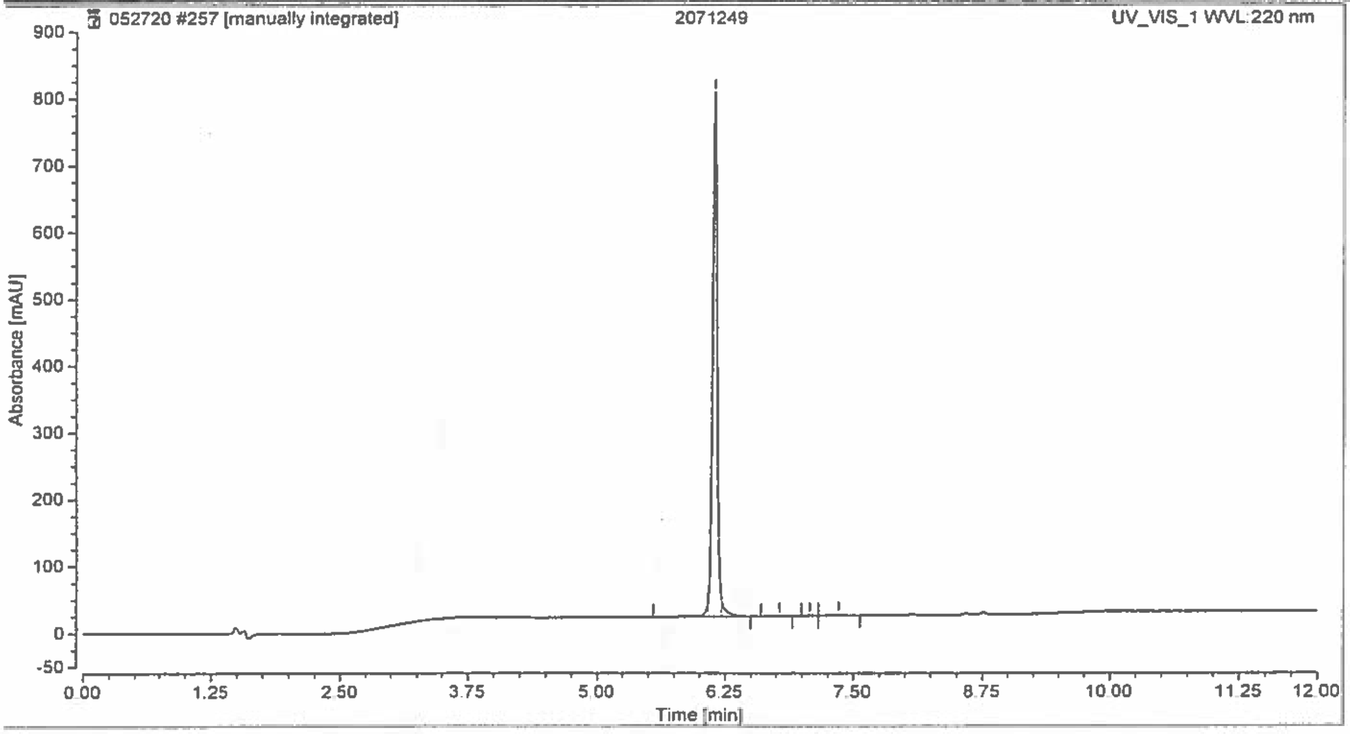

#### Integration Results

| No. | Retention Time min | Area mAU*min | Height mAU | Relative Area % |
|---|---|---|---|---|
| 1 | 6.083 | 0.670 | 19.269 | 1.55 |
| 2 | 6.147 | 41.100 | 784.336 | 95.32 |
| 3 | 6.217 | 1.101 | 25.057 | 2.55 |
| 4 | 6.783 | 0.024 | 0.553 | 0.06 |
| 5 | 7.073 | 0.036 | 0.654 | 0.08 |
| 6 | 7.360 | 0.186 | 1.409 | 0.43 |
| **Total:** | | **43.117** | **831.279** | **100.00** |

C:\Xcalibur\Data\2071249 06/18/20 15:05:23

06/19/20

2071249 #36-108 RT: 0.24-0.69 AV: 24 NL: 3.93E5
F: ITMS + c ESI E Full ms [50.00-2000.00]

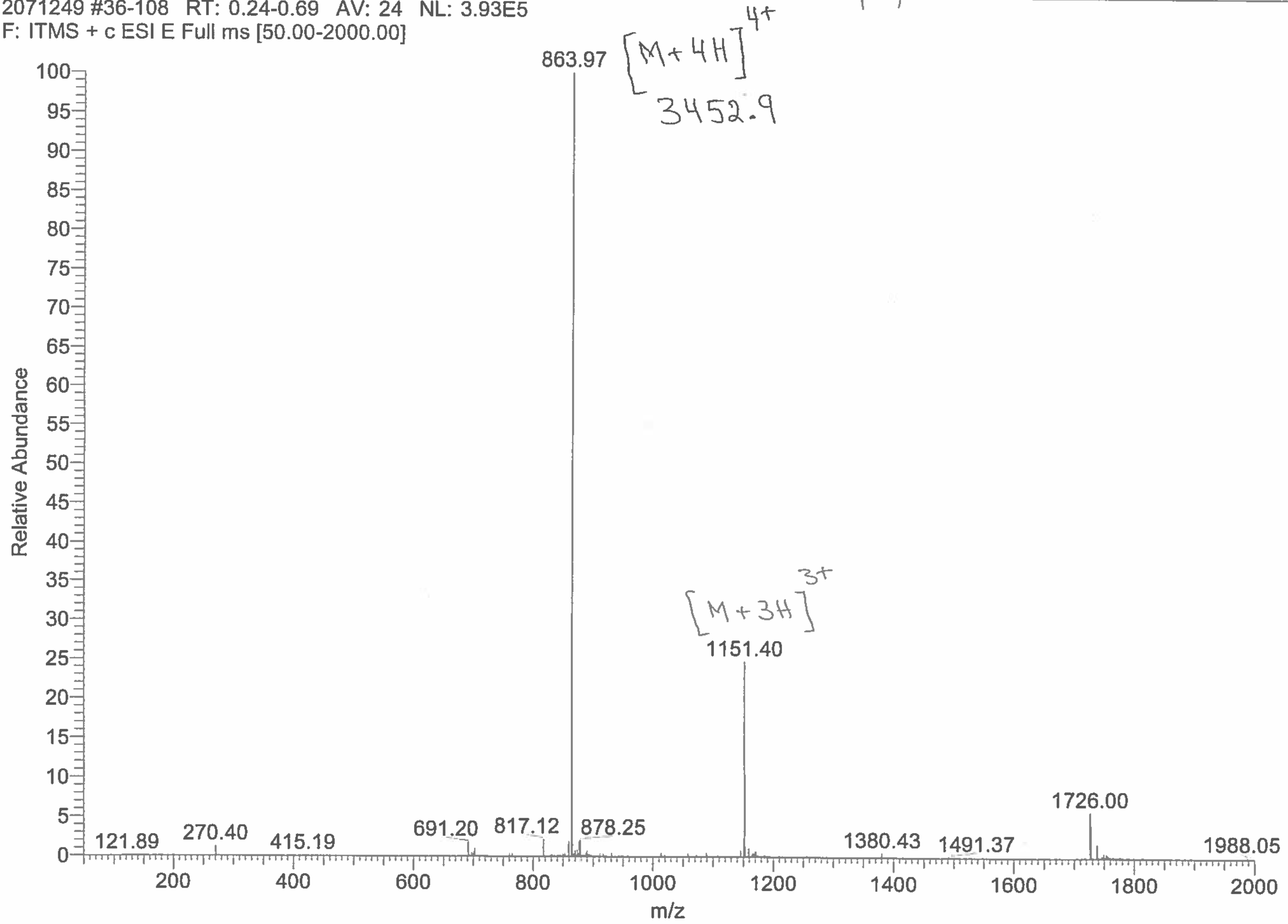

Eurogentec

### ANASPEC PEPTIDE
#### QC Data Sheet

ANASPEC

For Research Use Only

| YOUR ORDER | SON | DATE |
|---|---|---|
| B001890494 | 1500055456 | 19-Jun-20 |

| CUSTOMER | ADDRESS / INSTITUTION |
|---|---|
| University of California San Francisco (UCSF) | UNITED STATES San Francisco |

| PEPTIDE NAME | LOT# | INTERIM | SCALE | # AMINO |
|---|---|---|---|---|
| 75386-6: (disulfide bridge) | **2056280** | 2071250 | Custom | 26 |

**SEQUENCE (N-Term → C-Term)**

**NOTA -X-** DY SHC(S-) SPL RYY PWW KC(S-)T YPD PEG GG **- NH2** , with disulfide bridge
**X=AEEEA**

| PHYSICOCHEMICAL PROPERTIES | |
|---|---|
| 1A 280 [mg/ml] * | 0.2 |
| Charged at pH 7 * | 0.0 |
| Isoelectric Point * | 6.7 |

*(* Theoretical values)*

| REGULAR AA PROPERTIES | | | | | |
|---|---|---|---|---|---|
| Charged AA | D,H,R,K,E | 6 | Polar AA | D,S,H,R,K,T,E | 9 |
| Acid AA | D,E | 3 | Hydrophobic AA | Y,P,L,W,G | 1 |
| Basic AA | H,R,K | 3 | | | |

**QC DATA**

| Attribute | Test method | Acceptance criteria | Result |
|---|---|---|---|
| Appearance | Visual | Report result | White Powder |
| % Peak Area by HPLC | HPLC | ≥ 95 % | 95 % |
| Identity | MS | 3449.8 ± 0.2 % | 3451.0 |

**DELIVERABLE**

**Format** Dried

**Aliquoting**
Number of Aliquots 1
Qty by Aliquot (mg) 10 mg

LOT# 2056280 10 mg
(disulfide bride)
For Laboratory Use Only
ANASPEC

**DELIVERY CONDITION**
Room temperature

**STORAGE CONDITION**
-20 °C, dry

**COMMENTS**

**PEPTIDE RECONSTITUTION AND STORAGE**

Please read the entire section before proceeding with the solubilization of your custom peptide.

Peptides are shipped at ambient temperature as a lyophilized powder. Upon receipt store them at -20°C. Allow the vial to equilibrate to room temperature prior to opening.

Peptide solubility is highly dependent on the sequence. Peptides that are more hydrophobic (high propensity of A, F, G, V, L, I, M, W, P) in nature, will require an organic solvent in order to dissolve. Peptides that are acidic in nature (high propensity of D, E in the peptide sequence) require a basic aqueous buffer to dissolve, while peptides that are basic in nature (high propensity of K, H, and R) require an acidic aqueous buffer to dissolve.

To reconstitute a hydrophobic peptide, add 100 µL DMSO and sonicate until a homogenous solution forms. Next, add your buffer of choice to form a 1 mg/mL solution (a higher concentration of peptide will require a greater amount of DMSO). To reconstitute basic or acidic peptides, add 1 mL of the appropriate buffer to the peptide and sonicate to ensure a homogenous solution forms.

Reconstituted peptides can be stored frozen at -20°C for short period of time, but it is advisable to prepare multiple aliquots to avoid multiple freeze thaw cycles. We recommend that all aliquoted solutions be lyophilized if the peptide is going to be stored for extended periods of time at -20 °C.

Additionally, please note that peptides with a high propensity of basic residues (R, K, H) in their sequence may undergo a physical change from solid powder to an oil (via moisture absorption). This physical change does not affect the purity or functionality of the peptide.

Nomenclature used for the sequence termini:
- N-terminus: H means free amine ($NH_2$-), Ac mean acetyl [CH3C(O)-NH-], Pyr means pyroglutamic acid
- C-terminus: OH means free acid (-COOH), $NH_2$ means amide [-$CONH_2$]

Modifications on the side chain of amino acids are depicted in the parenthesis after the corresponding amino acid. For example; phosphorylated serine = S($PO_3H_2$) or epsilon-N-acetylated lysine = K(Ac)

**TECHNICAL SUPPORT**

If you have any questions feel free to call our Technical Support Centre

**EUROPE**
✆ 00 800 666 00 123 (European toll free number),
✉
Kaneka Eurogentec S.A. Liège Science Park
Rue Bois Saint-Jean 5 - 4102 SERAING BELGIUM
 Web: www.eurogentec.com
RPM Liège T.V.A.-(BE)-0427.348.346 - ING Belgique Bank - IBAN: BE86 3400 2118 6050 BIC: BBRUBEBB

**NORTH AMERICA**
✆ +1 800 452-5530 (American toll free number),
✉
AnaSpec, Inc. 34801 Campus Drive
Fremont, CA 94555 - USA
 Web: www.anaspec.com

### Chromatogram and Results

#### Injection Details

| | | | |
|---|---|---|---|
| *Injection Name:* | **2071250** | *Run Time (min):* | **12.50** |
| *Vial Number:* | **BA2** | *Injection Volume:* | **1.000** |
| *Column:* | **C18, 100 x 4.6mm, H18-074009** | *Channel:* | **UV_VIS_1** |
| *Instrument Method:* | **5-60%-7mins-Bextended-0.7ml-30C** | *Wavelength:* | **220.0** |
| *Processing Method:* | **test** | *Bandwidth:* | **4** |
| *Injection Date/Time* | **18/Jun/20 16:47** | *Instrument No.* | **QC-HPLC-9** |

#### Chromatogram

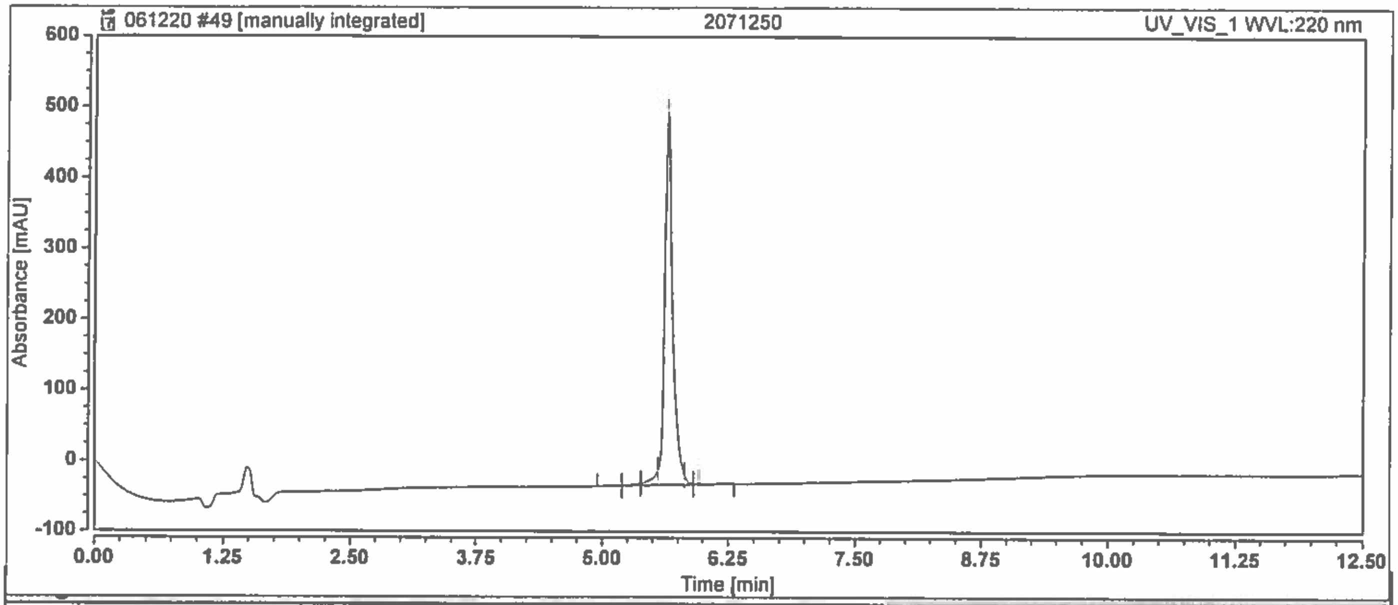

| No. | Retention Time min | Area mAU*min | Height mAU | Relative Area % |
|---|---|---|---|---|
| 1 | 5.197 | 0.014 | 0.000 | 0.03 |
| 2 | 5.380 | 0.288 | 3.131 | 0.58 |
| 3 | 5.553 | 1.483 | 21.673 | 2.97 |
| 4 | 5.633 | 47.498 | 526.989 | 95.25 |
| 5 | 5.813 | 0.411 | 13.648 | 0.82 |
| 6 | 5.960 | 0.171 | 2.507 | 0.34 |
| **Total:** | | **49.865** | **567.948** | **100.000** |

C:\Xcalibur\Data\2071250

06/18/20 15:08:09

06/19/20 [illegible]

2071250 #35-106 RT: 0.22-0.67 AV: 24 NL: 3.28E5
F: ITMS + c ESI E Full ms [50.00-2000.00]

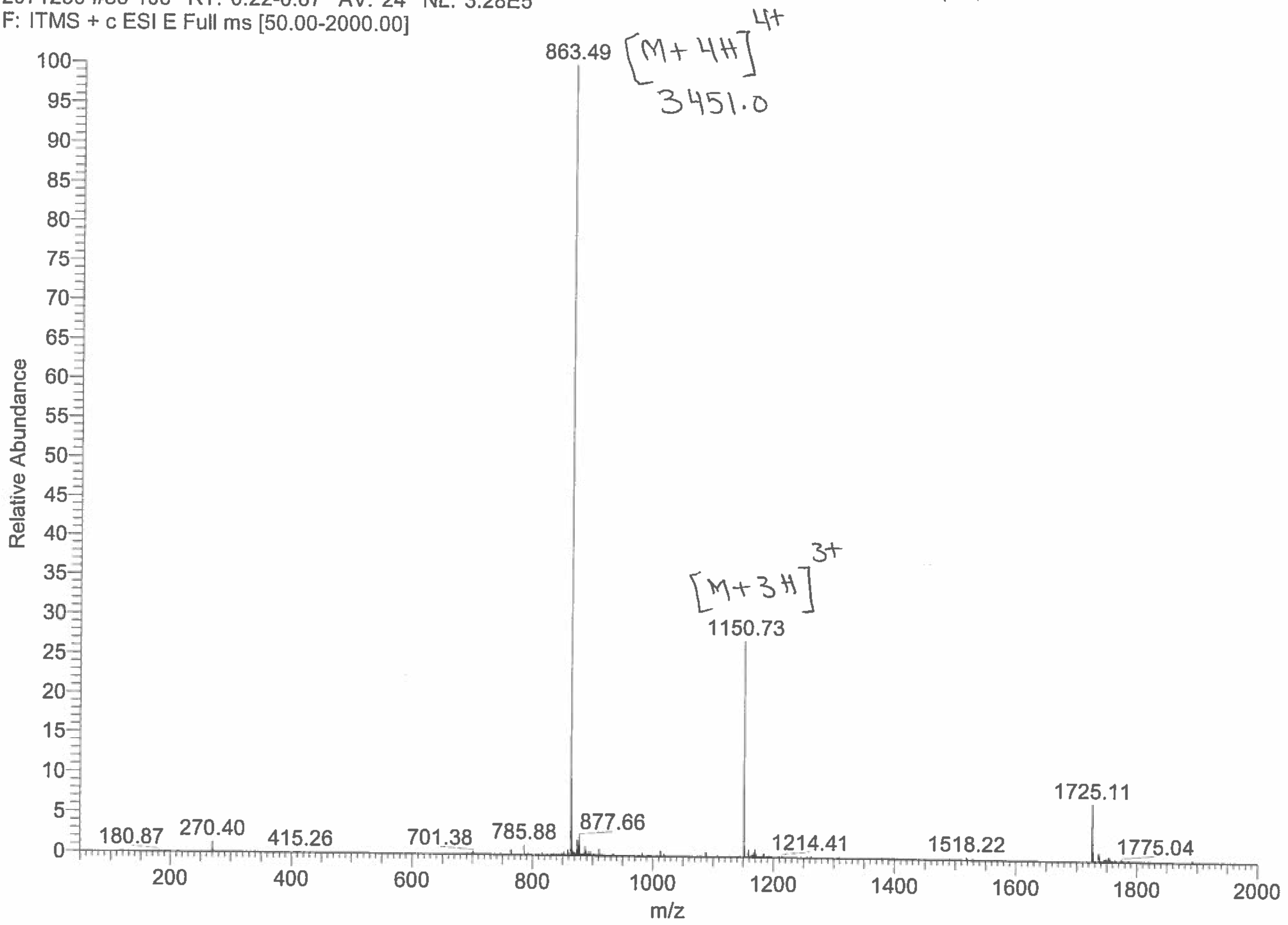
